# Human hnRNPA1 reorganizes telomere-bound Replication Protein A

**DOI:** 10.1101/2023.05.09.540056

**Authors:** Sophie L. Granger, Richa Sharma, Vikas Kaushik, Mortezaali Razzaghi, Masayoshi Honda, Paras Gaur, Divya S. Bhat, Sabryn M. Labenz, Jenna E. Heinen, Blaine A. Williams, S. M. Ali Tabei, Marcin W. Wlodarski, Edwin Antony, Maria Spies

**Affiliations:** Department of Biochemistry and Molecular Biology, University of Iowa Carver College of Medicine, 51 Newton Road, Iowa City, IA 52242, USA; Department of Hematology, St Jude Children’s Research Hospital, Memphis, TN 38105, USA; Department of Biochemistry and Molecular Biology, St. Louis University School of Medicine, 1250 Carr Lane, St. Louis, MO 63104, USA; Department of Physics, University of Northern Iowa, Cedar Falls, IA 50614, USA

**Keywords:** Telomere, telomere biology disorder, G-quadruplex, Replication Protein A (RPA), heterogeneous nuclear ribonucleoprotein A1 (hnRNPA1), telomeric repeat-containing RNA (TERRA), single-molecule total internal reflection fluorescence microscopy (smTIRFM), Forster resonance energy transfer (FRET), mass photometry (MP)

## Abstract

Human replication protein A (RPA) is a heterotrimeric ssDNA binding protein responsible for many aspects of cellular DNA metabolism. Dynamic interactions of the four RPA DNA binding domains (DBDs) with DNA control replacement of RPA by downstream proteins in various cellular metabolic pathways. RPA plays several important functions at telomeres where it binds to and melts telomeric G-quadruplexes, non-canonical DNA structures formed at the G-rich telomeric ssDNA overhangs. Here, we combine single-molecule total internal reflection fluorescence microscopy (smTIRFM) and mass photometry (MP) with biophysical and biochemical analyses to demonstrate that heterogeneous nuclear ribonucleoprotein A1 (hnRNPA1) specifically remodels RPA bound to telomeric ssDNA by dampening the RPA configurational dynamics and forming a ternary complex. Uniquely, among hnRNPA1 target RNAs, telomeric repeat-containing RNA (TERRA) is selectively capable of releasing hnRNPA1 from the RPA-telomeric DNA complex. We speculate that this telomere specific RPA-DNA-hnRNPA1 complex is an important structure in telomere protection.

**One Sentence Summary:** At the single-stranded ends of human telomeres, the heterogeneous nuclear ribonucleoprotein A1 (hnRNPA1) binds to and modulates conformational dynamics of the ssDNA binding protein RPA forming a ternary complex which is controlled by telomeric repeat-containing RNA (TERRA).

## Introduction

Replication protein A (RPA) coordinates a plethora of DNA metabolic events by binding to virtually all exposed single-strand (ss)DNA in the cell. RPA serves as an interaction hub that recruits over three dozen proteins onto ssDNA, melts secondary DNA structures, activates the DNA damage response, and hands off ssDNA to appropriate downstream proteins^1,2^. RPA functions as a stable heterotrimer composed of RPA1 (70 kDa), RPA2 (32 kDa) and RPA3 (14 kDa) subunits with modular oligosaccharide/oligonucleotide binding (OB) domains spread across RPA1 (OB-F, A, B and C), RPA2 (OB-D), and RPA3 (OB-E)^1^. Functionally, these OB-domains are further classified as DNA-binding domains (DBDs-A, B, C & D) and protein-interaction domains. Since these domains are connected by flexible linkers, the four DBDs allow for dynamic protein-ssDNA interactions, whereby a macroscopically bound RPA cycles between high affinity and low affinity binding modes, and thus can be displaced or remodeled by lower affinity DNA binding proteins acting downstream of RPA^1–6^. RPA binds ssDNA with high affinity (sub-nM Kd) and little sequence specificity, though it displays a preference for pyrimidines over purines^7^, as well as for G-rich^8^ and telomeric DNA^9^.

Human telomeres are nucleoprotein structures made of tandem (TTAGGG) repeats with single-strand G-rich overhangs and shelterin complex proteins^11,12^. Non-canonical DNA structures including G-quadruplexes formed by the telomere G-rich strand^13–15^ are enriched at telomeric ends, and can cause replication fork stalling and telomere replication stress,^16,17^ and contribute to telomere dysfunction, DNA damage response and accelerated aging^18,19^. RPA localization at telomeres has been observed during telomere replication^10^. RPA melts telomere G-quadruplexes to maintain replication and protects telomeric ssDNA from being recognized and targeted by DNA repair machinery^16^. In agreement with a delicate balance of RPA functions at telomeres, we have recently identified germline heterozygous *RPA1* dominant gain of function mutations in patients with telomere biology disorder, characterized by pathological shortening of telomeres resulting in bone marrow failure, liver and lung fibrosis, mucocutaneous fragility and predisposition to cancers^9^. Specifically, purified RPA carrying the p.E240K mutation in RPA1 (denoted as E240K moving forward) binds ssDNA with higher affinity than the wild type protein and displays enhanced capacity to melt telomeric G-quadruplexes.

While the role of RPA in sequestering and protecting ssDNA are well established, its function and regulation at telomeric ssDNA is poorly understood. RPA recruitment to the broken DNA with a single strand overhang ensures proper DNA repair. However, its recruitment to natural chromosome ends can result in deleterious end-to-end fusions. To protect chromosome ends, the shelterin complex binds telomeres, and protects telomeric ssDNA via the telomere-specific ssDNA-binding protein, Protection of Telomeres 1 (POT1). POT1 prevents RPA recruitment and subsequent ATR driven DNA damage response at telomeres. RPA transiently binds telomere overhangs during DNA replication in S-phase but must be quickly replaced by POT1 to ensure end protection. This process is tightly regulated by heterogeneous nuclear ribonucleoprotein A1 (hnRNPA1), an RNA binding protein that also binds chromosomal ends. Activities of hnRNPA1 at telomeres are in turn regulated by the telomeric repeat-containing RNA (TERRA), a long non-coding RNA transcribed from sub-telomeric regions towards chromosome ends^20^. Specifically, Flynn and colleagues proposed that hnRNPA1 displaces RPA from single-stranded telomeric DNA thus allowing for POT1 binding and telomere protection, and that this activity is inhibited by TERRA RNA in early S-phase when RPA is needed for telomere replication^21^.

Using single-molecule total internal reflection fluorescence microscopy (smTIRFM), mass-photometry, and biochemical studies, we aimed to determine how the hnRNPA1 and TERRA control access to RPA-bound telomeres. Two distinct smTIRFM approaches allowed us to visualize the contacts between the DBDs of RPA and DNA through RPA labeling with an environmentally sensitive fluorescent dye, and the DNA geometry through single-molecule FRET (smFRET) between the DNA linked fluorophores. Unexpectedly, hnRNPA1 was unable to directly compete with RPA for telomeric ssDNA binding. Instead, we observed formation of the ternary complex between RPA, telomeric ssDNA and hnRNPA1. In this complex the two proteins physically interact, and hnRNPA1 alters the contacts between DBDs of RPA and DNA and alters the RPA’s conformational dynamics. While TERRA was a poor competitor of hnRNPA1 pre-bound to telomeric ssDNA, it readily removed hnRNPA1 from the RPA-DNA-hnRNPA1 complex restoring the architecture of the RPA-telomeric DNA complex. The formation of the RPA-DNA-hnRNPA1 complex and its remodeling are specific to telomeres and TERRA. Notably, hnRNPA1 was unable to remodel RPA complexes on the G-quadruplex forming *BCL-2* promoter 1245 sequence, *cMYC* promoter Pu27 sequence, or poly-dT. Likewise, the HIV-1 Exon Splicing Silencer 3 Element RNA was unable to remove hnRNPA1 from the RPA-DNA-hnRNPA1 complex despite strong interaction between this RNA and hnRNPA1^22^. Collectively, our data point towards an intricate choreography of a dynamic complex between telomeric ssDNA, RPA, and hnRNPA1 which allows for melting of telomeric G-quadruplexes, and reorganization of the protein-ssDNA complex.

## Results

### A dynamic RPA-ssDNA complex is altered by a gain of function E240K mutant RPA

Previously, we speculated that the E240K mutation in the DBD-A of RPA1 extends the DNA binding site explaining tighter binding of this RPA variant to ssDNA^9^. We expected this extended interaction to alter the microscopic configurational dynamics of mutant RPA, specifically of its DBD-A. To visualize the dynamic interaction of human RPA with ssDNA we produced human RPA and RPA^E240K^ heterotrimers labeled with an environmentally sensitive fluorescent dye MB543 at DBD-A (RPA-DBD-A^MB543^) or DBD-D (RPA-DBD-A^MB543^)^23,24^ (**Supplementary Figure 1**). Upon binding to ssDNA, MB543-labeled RPA produces an increase in fluorescence^3,24,25^. RPA-DNA interaction was monitored using smTIRFM experiments, in which biotinylated ssDNA ((dT)_100_) was tethered to the TIRFM flow cell surface illuminated with 532 nm evanescent field. MB543-labeled protein was then flowed in, resulting in appearance of the fluorescent spots reflecting the presence of RPA on surface-tethered ssDNA (**Figure 1A&B**). MB543 proximity to DNA enhances this dye’s fluorescence yield^3,25^, with different modes of RPA-ssDNA interaction resulting in different levels of fluorescence (**Figure 1**). Global analysis of the fluorescence trajectories (time-based change in the fluorescence at specific locations on the slide) using a Bayesian inference-based approach implemented in hFRET software^26^ allowed us to assign the fluorescence states–from which we inferred the configurational states of the RPA-DNA complex. Our previous study of yeast RPA revealed several conformational states of the RPA-ssDNA complex with highest fluorescence level attributed to the most engaged state of the labeled domain^3^. Similar to its yeast counterpart, human RPA displays configurational dynamics when macroscopically bound to ssDNA (**Figure 1C, Supplementary Figure 2A&B**): both yeast and human RPAs labeled at DBD-A and DBD-D were found in four distinct configurational states characterized by different fluorescence intensities. The microscopic dynamics of human RPA DBD-A and DBD-D, however, was slower than that previously observed for yeast RPA^3^ with dwell times about 5-fold longer (**Figure 1C&D, Supplementary Figure 2**). RPA^E240K^-DBD-A^MB543^ displayed an unexpectedly complex fluorescence trend (**Figure 1E**). Its initial encounter with ssDNA resulted in a gradual increase in the MB543 fluorescence to levels much higher than those observed with wild type RPA-DBD-A^MB543^, RPA-DBD-D^MB543^, or mutant RPA^E240K^-DBD-D^MB543^ (see **Supplementary Figure 2** for examples of representative trajectories). After a period of 30-60 seconds, the MB543 fluorescence in the RPA^E240K^-DBD-A^MB543^ acquired step-like behavior similar to that observed for the wild type RPA-DBD-A^MB543^, though much brighter. This enhanced fluorescence may reflect the difference in the electrostatic environment of the dye brought about by the mutation and/or difference in the contacts between DBD-A and ssDNA due to the positively charged lysine in place of the negatively charged glutamic acid.

**Figure 1.**
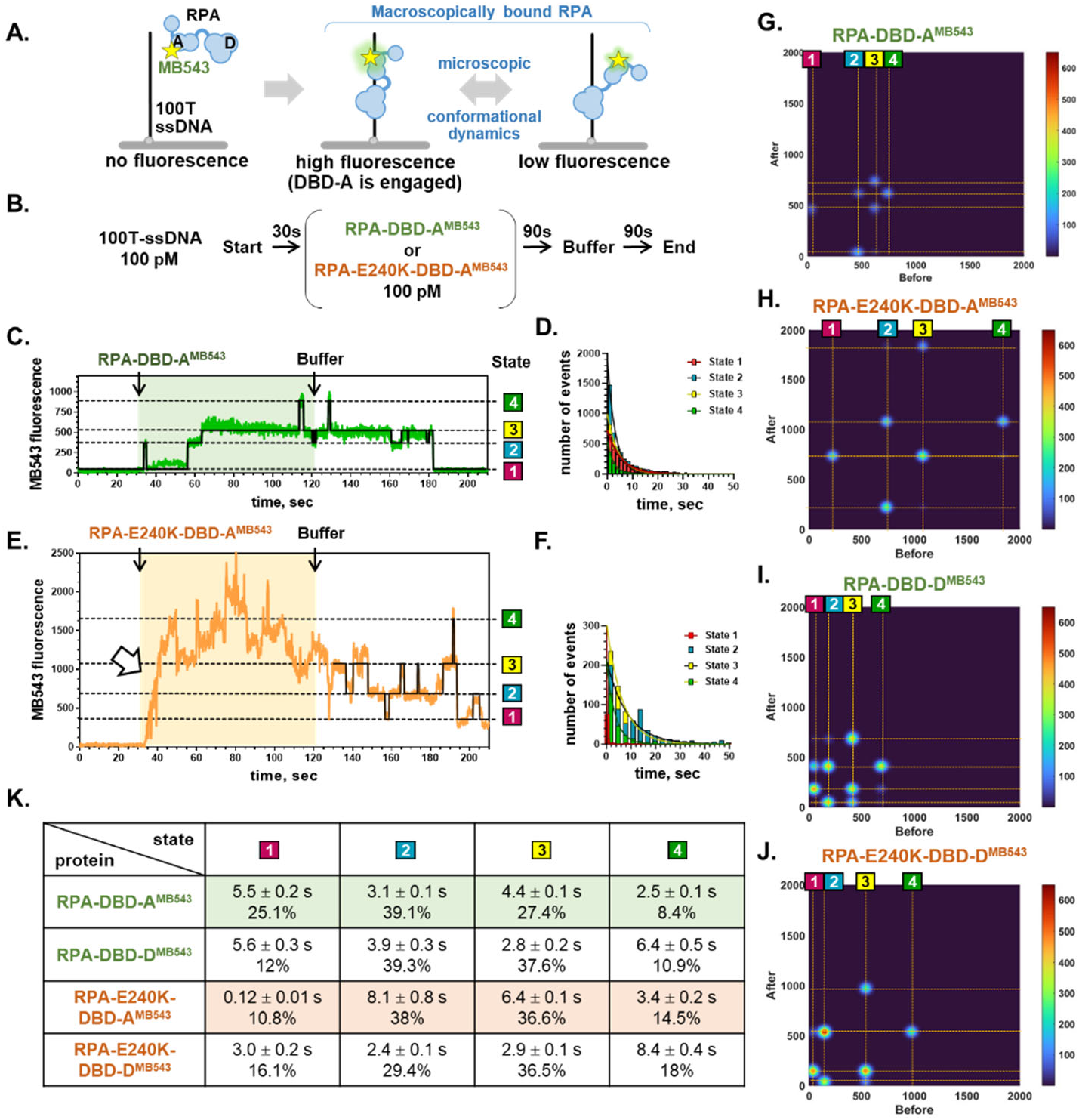
DBD-A of RPA^E240K^ displays altered conformational dynamics. **A.** Single-molecule TIRFM assay for monitoring the conformational dynamics of RPA. Biotinylated ssDNA molecules (dT100) are tethered to the surface of the smTIRFM flow cells. Binding of the MB543-labeled RPA manifests in the appearance of the fluorescence signal, while conformational dynamics of the DNA-bound RPA manifests in changes in the fluorescence intensity. **B**. Experimental scheme. **C**. A representative fluorescence trajectory (time-based changes in the MB543 fluorescence in a specific location in the smTIRFM flow cell) for the wild type RPA labeled with MB543 at the DBD-A (RPA-DBD-A^MB5^^43^; green) overlaid with an idealized trajectory (black) obtained by globally fitting all trajectories to a four-state model using hFRET. **D**. Dwell time distributions for the individual states of RPA-DBD-A^MB5^^43^. The dwell times were binned with a bin size of 2.4 seconds (bars). Solid lines represent exponential fits for each distribution. **E**. A representative fluorescence trajectory for the RPA^E240K^ mutant labeled with MB543 at the DBD-A (RPA^E240K^-DBD-A^MB5^^43^; orange) overlaid with an idealized trajectory (black). **F**. Dwell time distributions for the individual states of RPA^E240K^-DBD-A^MB5^^43^. The dwell times for states 2, 3 and 4 were binned with a bin size of 2.4 seconds (blue, yellow and green bars, respectively), while dwell times for state 1 were binned with a bin size 0.2 second (red bars). Solid lines represent exponential fits for each distribution. **G.-J.** Transition density plots for the RPA-DBD-A^MB5^^43^, RPA^E240K^-DBD-A^MB5^^43^, RPA-DBD-D^MB5^^43^, and RPA^E240K^-DBD-D^MB5^^43^, respectively. Only transitions after the buffer wash were considered in these plots. **K**. Summary of the dwell time analysis for all proteins in this study. The dwell times are shown as time constants from the exponential fitting of the dwell time distributions ± fitting error. Percentage of visitations to each state is shown below the respective dwell times. The total number of events used to build each distribution is shown in parentheses.

Our previous work showed that the stepwise changes in the yeast RPA-linked MB543 are due to configurational dynamics of the RPA-DNA complex and are not photophysical effects, and that the changes in the fluorescence can be attributed to a single RPA molecule^3^. Nevertheless, we only quantified the dwell times of the fluorescent states after removal of unbound RPA from the TIRFM flow cell. As evidenced by the transition density plots (**Figure 1G-J**), both RPA-DBD-A^MB5^^43^ and RPA^E240K^-DBD-A^MB5^^43^ progressed through their respective fluorescent and configurational states sequentially (1↔2↔3↔4), while the dynamics of the RPA-DBD-D^MB5^^43^ and RPA^E240K^-DBD-D^MB543^ included also 1↔3 and 2↔4 transitions. The dwell time distributions for all states were binned and fit with single exponential decay (see **Figure 1D&F** for the dwell-time distributions of the RPA-DBD-A^MB543^ and RPA^E240K^-DBD-A^MB543^, respectively) and are summarized in **Figure 1K**. The main difference between the two proteins was in the duration of the least engaged state (State 1) of the DBD-A, which was reduced in the RPA^E240K^-DBD-A^MB543^ to ∼0.12 seconds compared to ∼5.5 seconds in the wild type RPA-DBD-A^MB543^. Notably, both the initial gradual increase in the MB543 fluorescence and the dwell times in each state were independent of the RPA^E240K^-DBD-A^MB543^ concentration, while the number of observed trajectories and individual events increased linearly with increasing protein concentration (**Supplementary Figure 3**). This confirms that the observed phenomenon and the quantified dwell times reflect the behavior of RPA-DNA complexes containing a single RPA^E240K^-DBD-A^MB543^ protein. These data support our earlier model that the mutation drives an alternate configuration for DBD-A on ssDNA with an extended DNA binding site. It is important to note that we monitor the changes in configuration of the RPA-ssDNA complex associated with DBD-A, and it is very likely that DBD-B works in concert with DBD-A^27^. In contrast to the DBD-A, the conformational dynamics of the DBD-D was similar for the wild type and mutant protein (**Figure 1 and Supplementary Figure 2**).

### Improved association of RPA^E240K^ with telomeric G-quadruplex

Human RPA has a capacity to melt DNA secondary structures including G-quadruplexes and this activity is enhanced by the E240K mutation^9,13–15^. While the efficient telomeric G-quadruplex unfolding was observed with nearly stoichiometric ratio of RPA and h-telG4 in solution studies using 10 nM DNA, sub-nanomolar concentrations were used in these smTIRFM experiments. Under such conditions RPA transiently binds telomeric G-quadruplex folded in the presence of potassium, and, when bound, RPA-DBD-A^MB543^ mostly spends time in the low fluorescence state (**Figure 2, Supplementary Figure 4**). However, we observed high fluorescent states in the beginning of most of the individual trajectories (**Figure 2, Supplementary Figure 4**). Both the wild type RPA and RPA^E240K^ displayed dynamic changes when added to the reaction chamber with immobilized human telomeric G-quadruplex DNA (five ATTGGG repeats and a dsDNA region used for DNA tethering; **Figure 2A**). These changes in fluorescence may be attributed to configurational dynamics of individual RPAs or to simultaneous binding of multiple RPAs. Notably, virtually all trajectories for the wild type RPA displayed protein dissociation or transition into state 1 with rare excursions to state 2 when excess of RPA was removed from the reaction chamber. In contrast, RPA^E240K^-DBD-A^MB543^ continued displaying changes in fluorescence upon removal of unbound protein consistent with enhanced stability of its G-quadruplex binding.

**Figure 2.**
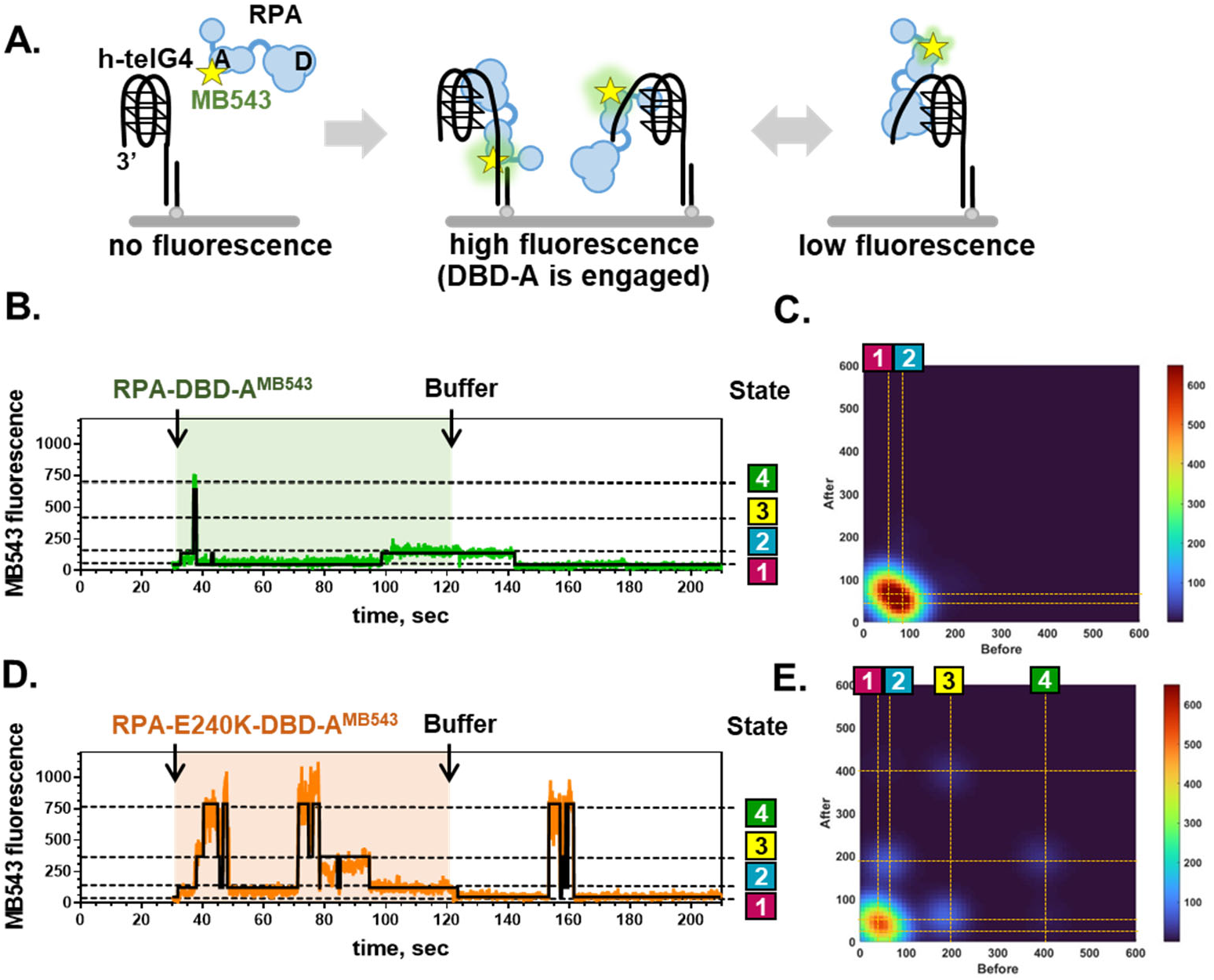
Binding and conformational dynamics of RPA-DBD-A^MB543^ and RPA^E240K^-DBD-A^MB543^ on telomeric G-quadruplex DNA. **A.** Experimental setup. Biotinylated partial duplex DNA with 3’-ssDNA overhang containing five TTAGGG repeats was prepared under conditions enforcing formation of the G-quadruplex, and tethered to the TIRFM flow cell. At 30 seconds, the indicated fluorescently-labeled RPA was flowed in, and at 120 seconds the unbound protein was removed by flowing in buffer. **B-E**. Representative MB543 trajectories and transition density plots (TDPs) for RPA-DBD-A^MB5^^43^ (**B&C**) and RPA^E240K^-DBD-A^MB5^^43^ (**D&E**), respectively. TDPs were built only from the events after buffer wash.

Discrepancy between inability of individual RPA molecules to unfold h-telG4 DNA in single-molecule experiments^13,15^ where 100 pM DNA was tethered to the surface, and bulk experiments carried out in the presence of 10 nM DNA, prompted us to identify conditions where we can observe individual RPA molecules stably bound to h-telG4 DNA under single-molecule conditions. Equilibrium smFRET titrations (**Supplementary Figure 5A**) showed that while mid to high nM concentrations of RPA were required to extend surface-tethered h-telG4 DNA in K^+^-containing buffer (**Supplementary Figures 5B and 6C**), stoichiometric DNA-RPA complexes were formed and were sufficient to extend the h-telG4 DNA in Li^+^-containing buffer (**Supplementary Figures 5C and 6D**). Therefore, in the experiments below we monitored configurational RPA dynamics on telomeric ssDNA in the Li^+^-containing buffers.

### RPA and hnRNPA1 physically interact and form ternary complexes on telomeric ssDNA *in vitro* and in cells

Human hnRNPA1 was reported to act as a key mediator of the RPA to POT1 exchange^21^. It interacts with both telomeric ssDNA and TERRA RNA using two RNA binding domains which comprise a structured UP1 region, as well as a RGG box in the unstructured C-terminal region^28–31^. To assess the effect of hnRNPA1 on the RPA-telomeric DNA complex we first carried out bulk FRET-based experiments that followed the geometry of the ssDNA in complex with RPA, hnRNPA1, and RPA plus hnRNPA1 combined (**Figure 3**). The unstructured ssDNA is extended to nearly contour length when bound by RPA, which can be visualized by incorporating FRET donor (Cy3) and FRET acceptor (Cy5) at the ends of the ssDNA and following change in FRET upon RPA binding (0.22 FRET)^32,33^. Similarly, G-quadruplex unfolding by RPA yields a FRET signal (0.59 in K^+^ or 0.55 in Li^+^) consistent with partially extended telomeric ssDNA or an equilibrium between folded and unfolded G-quadruplex^9^.

**Figure 3.**
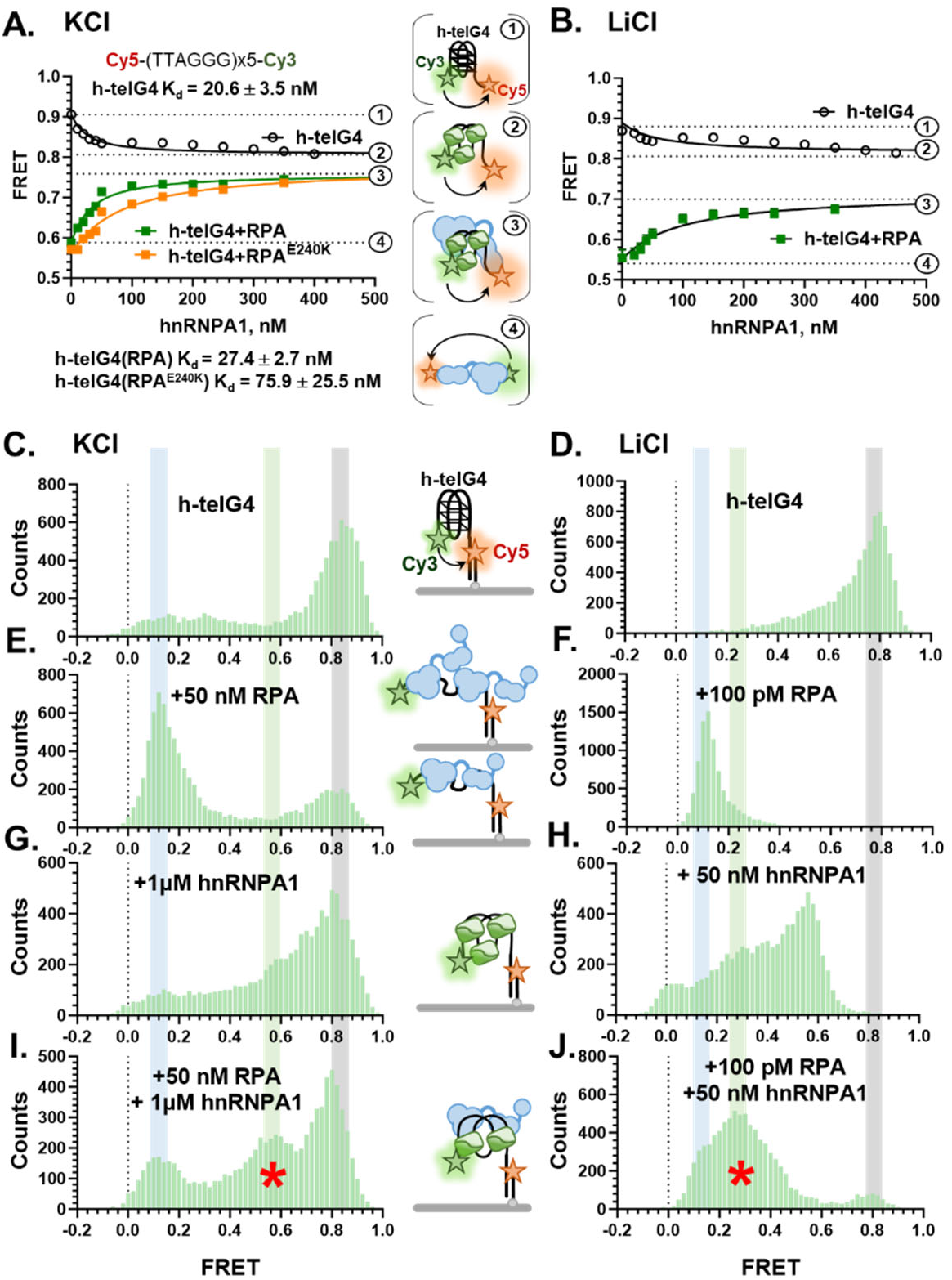
RPA and hnRNPA1 reorganize telomeric G-quadruplex DNA and form ternary complex. **A.** Bulk FRET-based analysis of the hnRNPA1 binding to 10 nM telomeric G-quadruplex folded in K^+^ containing buffer (open black circles), and to telomeric G-quadruplex melted by RPA (green squares) or RPA^E240K^ (orange squares). The binding curves were fitted using quadratic binding equation, and the K_d_s are shown with their respective fitting errors. Each complex with their respective FRET values are shown schematically on the right labeled with Cy3 (FRET donor) and Cy5 (FRET acceptor) at the two termini. The free and bound DNA substrates are schematically shown on the right with their respective FRET values. The experiments were carried out in triplicates and the values are plotted as average ± standard deviation for the three independent titrations. Note that the error bars for most measurements were smaller than the symbols’ sizes. Apparent equilibrium dissociation constants are shown with their respective fitting errors. **B.** hnRNPA1 binding to 10 nM telomeric G-quadruplex folded in Li^+^ containing buffer (open black circles), and to telomeric G-quadruplex melted by RPA (green squares). **C-J**. Equilibrium smFRET distributions for the protein-free h-telG4 labeled with the Cy3 and Cy5 dyes in K^+^ (**C**) and Li^+^ containing buffer (**D**), h-telG4 bound by saturating amounts of RPA (**E&F**), hnRNPA1 (**G&H**), and RPA and hnRNPA1 together (**I&J**). Green bars and red asterisks mark FRET species specific to the h-telG4/RPA/hnRNPA1 complexes. For all measurements, FRET values collected from 10 randomly chosen fields of view, binned in 0.02 unit bins and plotted as histograms. Experiments were repeated at least three times. A representative set of distributions is shown.

Structural^30^ and single-molecule analyses^34^ suggested that hnRNPA1 binding to human telomeric ssDNA reorganizes the quadruplex by juxtaposing the individual telomeric repeats. Using bulk FRET-based assays, we confirmed that purified hnRNPA1 preferentially binds telomeric ssDNA with an apparent K_d_ = 73.5±4.3 nM (**Supplementary Figure 7**). These assays utilized Cy3/Cy5-labeled, 15 nucleotide long ssDNA containing 2.5 telomeric repeats, and therefore were too short to fold into a G-quadruplex. The high initial FRET (0.92) for 15-mer substrates was due to high compaction of the free ssDNA under our experimental conditions (150 mM K^+^ and 5 mM Mg^2+^). FRET values at the saturating hnRNPA1 concentrations (0.74) were consistent with the hnRNPA1 bridging the repeats^30^. For reference, RPA binding to and extension of 15-mer telomeric DNA yields FRET of 0.6, and the RPA-mediated extension of dT15 ssDNA yields FRET of 0.42. Telomeric G-quadruplex formed by folding the Cy3/Cy5-labeled ssDNA containing five telomeric repeats in K^+^-containing buffer was bound by hnRNPA1 with apparent K_d_ = 20.6±3.5 nM and FRET of 0.81 at saturating hnRNPA1 concentrations (**Supplementary Figure 7**). The five-repeat long telomeric ssDNA (30 nucleotide) was used in these experiments because it corresponds to the binding site of one RPA heterotrimer. When presented with a stoichiometric complex of RPA and telomeric ssDNA (**Figure 3A**, green curve), hnRNPA1 binds to this complex with affinity similar to that of the telomeric G-quadruplex (apparent K_d_ = 27.4±2.7 nM). The saturating FRET value (0.76) is distinct from that of the telomeric G-quadruplex in complex with hnRNPA1 (0.81) suggesting that RPA is not fully displaced from the complex. Similar FRET values were observed when hnRNPA1 was titrated into nucleoprotein and Li complex containing RPA^E240K^ (**Figure 3A**, orange curve). The affinity of hnRNPA1 for RPA^E240K^-telomeric DNA complex was ∼4-fold lower (apparent K_d_ = 75.9±25.5 nM) than that for the complex containing wild type protein suggesting potential competition between the telomeric DNA engagement by the DBD-A of RPA and hnRNPA1. Similarly, hnRNPA1 binding to protein-free and RPA-bound htel-G4 in Li^+^-containing buffer results in distinct saturating bulk FRET values (**Figure 3B**).

In the smFRET experiments (**Figure 3C-J, Supplementary Figure 6**) we observed broad FRET distributions and the DNA-RPA-hnRNPA1 complexes with dynamic transitions between different FRET states. Unique FRET peaks, however, were readily observed in both K^+^ and Li^+^ conditions in the presence of both RPA and hnRNPA1 (**Figure 3I&J**, see peaks marked by a red asterisk). Formation of the ternary complex was further confirmed by EMSA experiments (**Supplementary Figure 8**). Complex between hnRNPA1 and human telomeric G4-quadruplex (h-telG4) DNA was readily detected with two to four molecules of hnRNPA1 being sufficient to shift all h-telG4 (**Supplementary Figure 8A**). When hnRNPA1 was added to the h-telG4 bound RPA, we observed a supershift indicative of the stable ternary complex formation (**Supplementary Figure 8B**). Notably, at high concentrations, hnRNPA1 was also able to bind unstructured ssDNA (**Supplementary Figure 8A**), however, no formation of the ternary complex was detected (**Supplementary Figure 8B**).

Our bulk and smFRET-based and EMSA-based analyses pointed towards coexistence of hnRNPA1 and RPA on the same DNA molecule contradictory to the previously suggested competition model^21^. To directly evaluate the molecular composition of the RPA-DNA-hnRNPA1 complex we used mass photometry (MP), a label-free single-molecule technique that applies interferometry to determine molecular mass of nucleoprotein complexes^35,36^ (**Figure 4A-J**, **Supplementary Table 2**). Both, RPA and RPA^E240K^ produced single Gaussian peaks with molecular weighs corresponding to an intact RPA heterotrimer (**Figure 4A&B**), while hnRNPA1 (38.7 kDa) appears as a peak with a mean molecular weight of 49.3 ± 0.1 kDa (S.D 10.9 ± 0.1 kDa) indicative of a mixture of monomers and dimers in solution (**Figure 4C**). Addition of the telomeric G-quadruplex DNA (five TTAGGG repeats) increased molecular weight of the RPA and RPA^E240K^ by 8 kDa (**Figure 4D&E**), while heterologous hnRNPA1-telomeric DNA complexes were observed (**Figure 4F**). To separate the effects of DNA binding and G-quadruplex unfolding, the G-quadruplex structure was destabilized by the presence of Li^+^ in the buffer. When both, RPA (or RPA^E240K^) and hnRNPA1 were mixed with the telomeric ssDNA, we observed an additional peak corresponding to the complex containing one DNA molecule, one RPA (or RPA^E240K^) heterotrimer and several hnRNPA1 molecules (**Figure 4G&H**). In the absence of DNA, we observed some complex formation between RPA and hnRNPA1 (**Figure 4I**), while added together, RPA^E240K^ and hnRNPA1 yielded two peaks corresponding to the two proteins (**Figure 4J**). To confirm that the RPA-hnRNPA1 complex was formed in cells we carried out co-immunoprecipitation assays in induced pluripotent stem cell (iPSC) lines that have either wild-type RPA1 or RPA^E240K^ (**Figure 4K**). Notably, both wild type RPA and RPA^E240K^ were found in the hnRNPA1 pull-down. RPA and hnRNPA1 are highly abundant nuclear proteins. Our ability to detect RPA^E240K^ in the hnRNPA1 pull-down, but not in the MP measurements likely reflects an IP of a DNA-mediated complex, as the RPA-hnRNPA1 or RPA^E240K^-hnRNPA1 complex may protect telomeric ssDNA from micrococcal nuclease digestion. While this work was in revision, BioID experiments confirmed that RPA and hnRNPA1 interact in the cell or are found in a close proximity to one another^37^.

**Figure 4.**
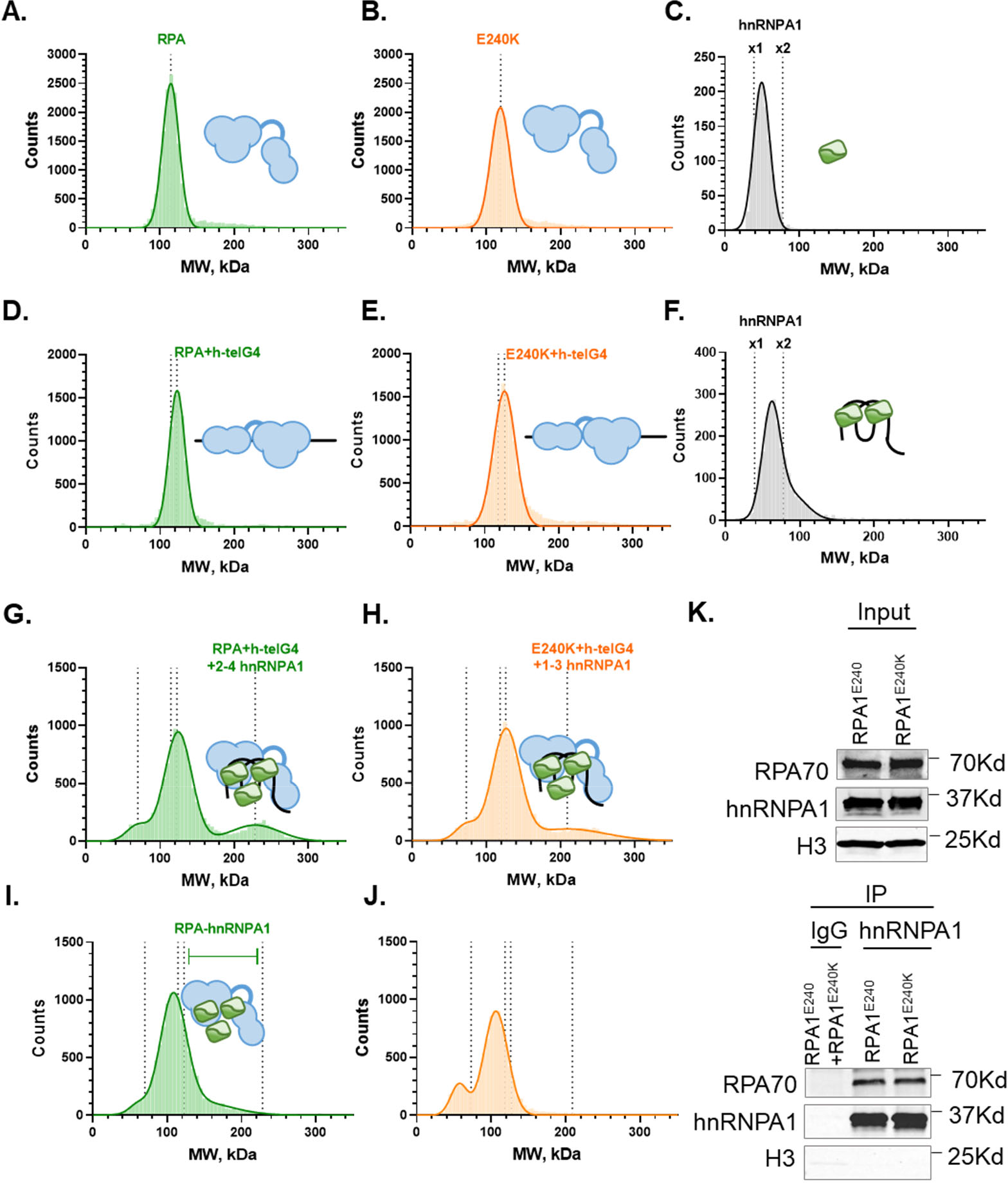
RPA and hnRNPA1 form ternary complex on telomeric ssDNA. **A-J.** The MP analyses of the 50 nM RPA (green), 50 nM RPA^E240K^ (orange) and 300 nM hnRNPA1 (grey), and their complexes with 50 nM telomeric ssDNA (five TTAGGG repeats) and each other in the Li^+^ containing buffer. Recorded mass values were binned in 3 kDa bins, plotted and fitted with one, two or three Gaussians (see Supplementary Table 2 for details). The optimal number of Gaussians used in each fit was selected based on an F-test. **A-C**. Individual proteins. **D-F.** Complexes of individual proteins and telomeric DNA. **G&H**. Molecular weights of RPA (**G**) or RPA^E240K^ (**H**) mixed with telomeric DNA and hnRNPA1. The new Gaussian peak which we attribute to the ternary RPA-DNA-hnRNPA1 complex is marked with a cartoon representation of the complex. **I&J**. Molecular weights of RPA (**I**) or RPA^E240K^ (**J**) mixed with hnRNPA1. **K.** hnRNPA1 physically associates with RPA in cell lysates. Cell lysates of PCS201 iPSCs with wild type RPA1 and RPA1 p.E240K were treated with micrococcal nuclease prior to immunoprecipitation with anti-hnRNPA1. *N* = 3 biologically independent experiments. Histone H3 was used as a loading control.

### hnRNPA1 constrains the configurational dynamics of RPA on telomeric DNA

When RPA-DBD-A^MB543^ or RPA^E240K^-DBD-A^MB543^ was bound to individual DNA molecules containing 3′-ssDNA overhang with five telomeric repeats in G-quadruplex destabilizing buffer containing Li^+^, we observed a dynamic behavior similar to that observed on the (dT)_100_ ssDNA (**Figure 5 and Supplementary Figure 9**). Addition of hnRNPA1 caused gradual decrease in fluorescence that stabilized at a lower fluorescence for the wild type RPA (**Figure 5C and Supplementary Figure 9A**), and at an intermediate fluorescence for RPA^E240K^ (**Figure 5D and Supplementary Figure 9B**), suggesting that first, the RPA dynamics is constrained in the RPA-DNA-hnRNPA1 complex, and second, that DBD-A is not fully engaging the DNA in this complex. A rarer excursions from the low fluorescence state of RPA-DBD-A^MB543^ suggested that the RPA is still present in the complex. These observations were in complete agreement with the FRET and MP data demonstrated above.

**Figure 5.**
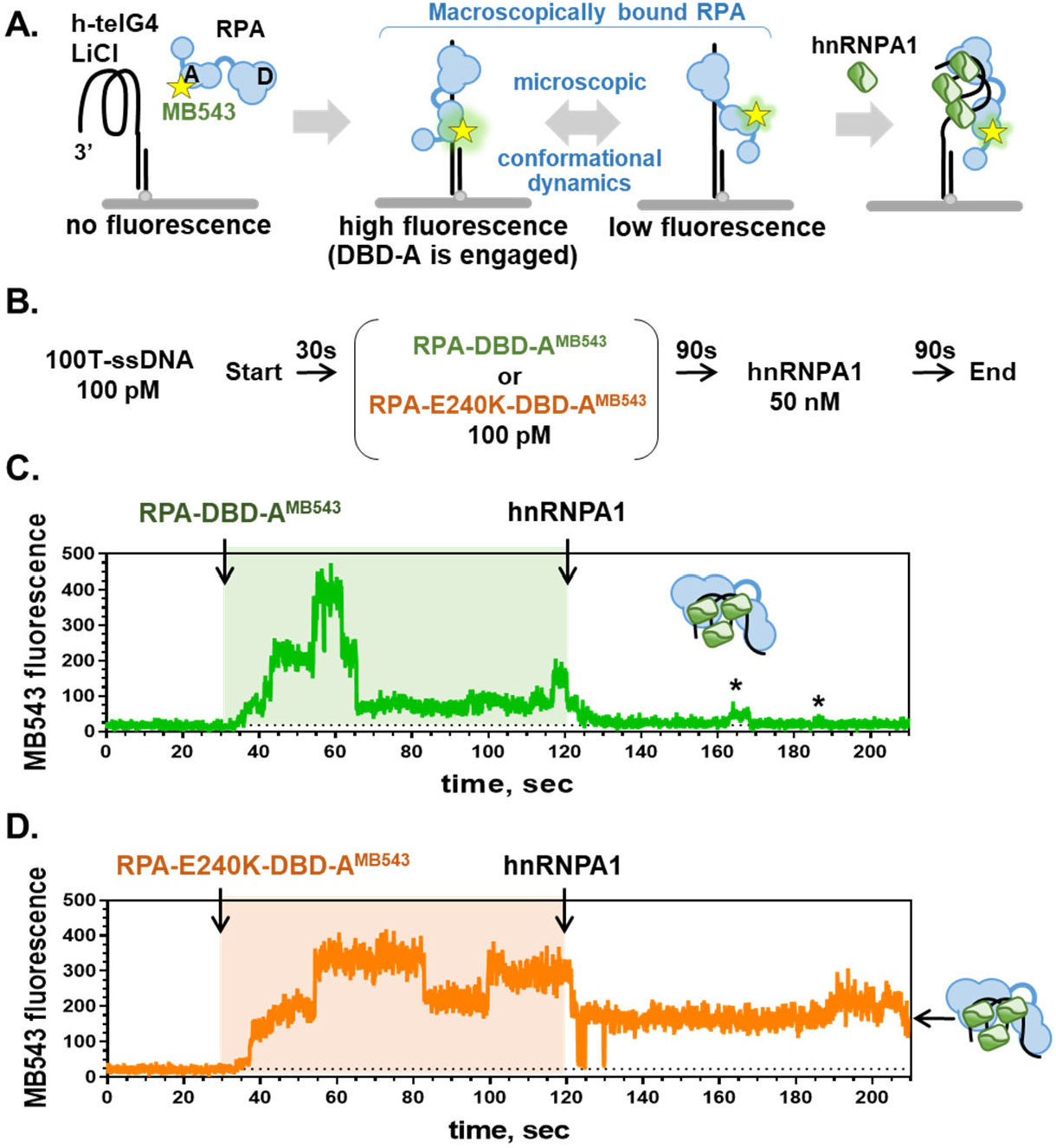
hnRNPA1 constrains conformational dynamics of the DBD-A of RPA bound to telomeric ssDNA. **A.** Experimental setup of the smTIRFM experiment. **B.** Experimental scheme. Biotinylated partial duplex DNA with 3’-ssDNA overhang containing five TTAGGG repeats was prepared in the Li^+^-containing buffer, and tethered to the TIRFM flow cell. At 30 seconds, the indicated fluorescently-labeled RPA was flowed in, and at 120 seconds the unbound protein was replaced with 50 nM unlabeled hnRNPA1. **C**. Representative MB543 trajectory for the wild type RPA-DBD-A^MB5^^43^. Asterisks mark deviation of the fluorescence signal from the lowest state suggestive of the presence of RPA. **D.** Representative MB543 trajectory for the RPA^E240K^-DBD-A^MB5^^43^. Signal level corresponding to the RPA^E240K^-hnRNPA1-DNA ternary complex is marked by an arrow and a carton representation of the complex.

### TERRA RNA controls the RPA-ssDNA-hnRNPA1 complex

We next set to determine how the presence of TERRA RNA affects the RPA-DNA-hnRNPA1 complex. To determine whether the same RPA molecule remains associated with telomeric ssDNA during hnRNPA1 binding and whether TERRA can remove the hnRNPA1 from the ternary complex, we carried out the following smTIRFM experiments (**Figure 6B**). A biotinylated DNA construct containing five repeats of human telomeric DNA was tethered on the surface of the TIRFM flow cell. RPA-DBD-A^MB543^ or RPA^E240K^-DBD-A^MB543^ (100 pM) was added, followed by the addition of 50 nM hnRNPA1, and then 50 nM (molecules) TERRA RNA. After addition of each component, the flow cell was allowed to equilibrate for five minutes and five short movies (100 frames) were recorded in different regions of the flow cell. The flow cell was illuminated only during recording of the movies to reduce photobleaching. Representative frames shown in **Figure 6B** highlight the appearance of the fluorescent signal upon addition of RPA-DBD-A^MB543^ (dark spots). Fainter fluorescence spots are observed upon addition of hnRNPA1 to the RPA-DNA complex, but the signals recover after addition of TERRA indicating that the RPA-DBD-A^MB543^ remained associated with surface-tethered DNA after addition of hnRNPA1, but its DBD-A was constrained in a dark/less engaged state.

**Figure 6.**
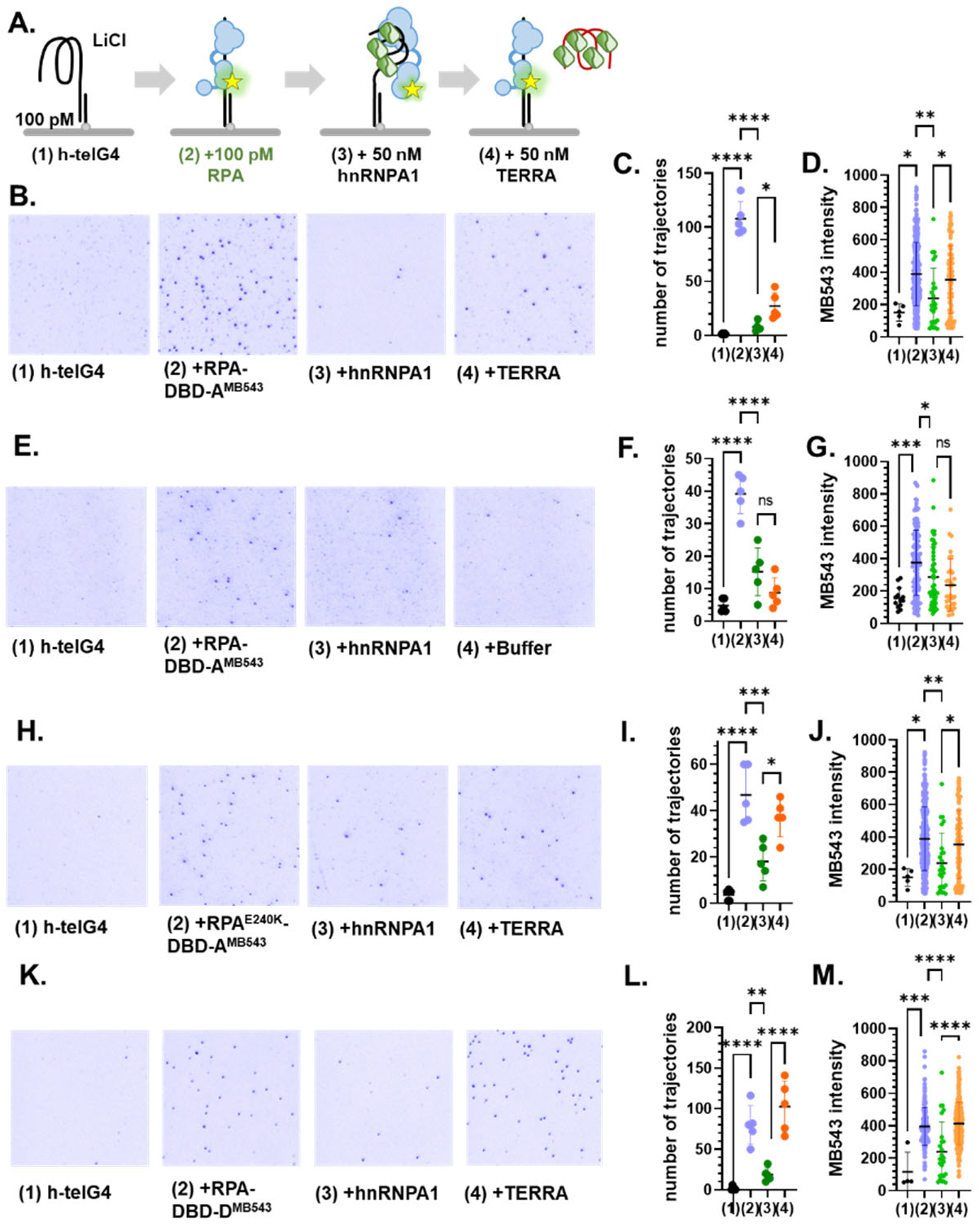
RPA-hnRNPA1 complex on telomeric ssDNA is remodeled by TERRA RNA. **A.** Experimental scheme. Short movies were recorded under equilibrium conditions in the Li^+^-containing buffer (1) in the presence of tethered DNA, (2) upon addition of 100 pM of indicated fluorescently labeled RPA, (3) upon replacement of unbound RPA with 50 nM hnRNPA1, and (4) upon replacement of hnRNPA1 with buffer or 50 nM (molecules) TERRA RNA. Representative color-inverted images of the 1/4 of the field of view are shown in panels **B**, **E**, **H**, and **K**. Number of trajectories with the fluorescence signal above background recorded in five short movies under each conditions are quantified in panels **C**, **F**, **I**, and **L**. Fluorescence in each point of these trajectories is summarized in panels **D**, **G**, **J**, and **M**. Statistical analysis: ordinary one-way ANOVA (GraphPad Prism).

The data from five replicates for each condition were quantified to reveal the number of observed trajectories per movie (**Figure 6C**), and fluorescence intensity within the selected trajectories across 100 frames of each movie (**Figure 6D**). Addition of TERRA resulted in recovery of the trajectories that show fluorescence that can be distinguished form the background and overall fluorescence intensity in these trajectories. No fluorescence above background was observed in control experiments without MB543-labeled RPA. When the excess of hnRNPA1 was removed by addition of buffer instead of TERRA, we did not observe the RPA-DBD-A^MB543^ fluorescence recovery, confirming stability of the RPA-DNA-hnRNPA1 complex (**Figure 6E-G**). Similar to the wild type protein, fluorescence of the RPA^E240K^-DBD-A^MB543^ was reduced after addition of hnRNPA1 and then recovered upon addition of TERRA (**Figure 6H-J**). Similar behavior was observed with RPA-DBD-D^MB543^ (**Figure 6K-M**) suggesting that hnRNPA1 remodels and constrains the whole RPA.

**Supplementary Figure S10** shows bulk FRET-based experiments where we pre-incubated the Cy3/Cy5-labeled telomeric G-quadruplex (five telomeric repeats) folded in the presence of K^+^ with RPA, hnRNPA1, or both proteins. We then titrated unlabeled TERRA RNA (three UUAGGG repeats) into these complexes. TERRA had no effect on the RPA bound to telomeric DNA (filled green circles). Unexpectedly, TERRA also had little effect on the pre-formed complex between hnRNPA1 and telomeric DNA in a broad range of TERRA concentrations (open black circles). However, when hnRNPA1 was titrated into the mixture of 10 nM Cy3/Cy5-labeled telomeric DNA and 100 nM TERRA RNA, higher hnRNPA1 concentrations were needed to achieve G-quadruplex binding, confirming competition between TERRA and telomeric DNA (not shown). In contrast, when TERRA was titrated into the pre-formed RPA-DNA-hnRNPA1 complex we observed hnRNPA1 release and return of the FRET signal to the level corresponding to RPA-DNA complex with an apparent K_d_ = 8.2±0.7 nM.

### Formation of the RPA-DNA-hnRNPA1 complex is specific for telomeric ssDNA

To probe whether the hnRNPA1 remodeling of RPA-ssDNA complex is telomere-specific, we performed FRET-based and EMSA experiments on three additional DNA substrates decorated at termini similar to the human telomeric G-quadruplex with the Cy3 and Cy5 dyes, (dT)_30_, *BCL-2* promoter 1245 G-quadruplex^38^, and *cMYC* promoter Pu27 G-quadruplex^39^. While RPA was able to bind and form a stoichiometric 1:1 complex with each of these substrates (**Supplementary Figure 11**), the FRET level at saturating RPA concentrations was different for the different DNAs suggesting a either difference in configuration between these complexes or different occupancy (**Supplementary Figure 11**). In contrast to telomeric G-quadruplex, however, hnRNPA1 was unable to remodel non-telomeric ssDNA structures or their complexes with RPA. These FRET-based analyses were further confirmed by the orthogonal EMSA experiments (**Supplementary Figure 12**). Both RPA and hnRNPA1 were able to bind BCL-2 and Pu27 G-quadruplexes, albeit less robustly compared to telomeric G-quadruplex. Complexes containing both proteins were readily observed on telomeric G-quadruplex, but only detectable at very high hnRNPA1 concentrations on BCL-2 and Pu27 G-quadruplexes.

### Telomere specific RPA-ssDNA-hnRNPA1 complex is not sensitive to non-telomeric hnRNPA1-interacting RNAs

Human hnRNPA1 binds many different cellular and viral RNAs, affects gene expression, RNA splicing, and viral infection. Earlier SELEX experiments identified a preferred RNA sequence, which contained a telomeric-like signature^40^. One of the viral RNA sequences recognized by hnRNPA1 is the HIV viral splicing silencer (ESS3), which is bound by hnRNPA1 with similar affinity to TERRA RNA^22^. To test whether control of the RPA-DNA-hnRNPA1 complex by TERRA RNA is unique to its sequence we have compared the effects of TERRA, SELEX-derived RNA, and HIV ESS3 RNA on the conformation of the telomeric ssDNA bound by RPA and hnRNPA1 (**Supplementary Figure 11B**). Similar to TERRA, SELEX-derived RNA was able to remove hnRNPA1 from the complex. HIV ESS3 RNA on the other hand had little effect on the configuration of the telomeric DNA bound by RPA and hnRNPA1 suggesting that in its presence, hnRNPA1 remains in the complex.

## Discussion

Five to fifteen kilobases of repetitive telomeric DNA exists at the ends of chromosomes in a double stranded form with a 50-500 nucleotide G-strand overhang. Several molecular events at telomeres may provide opportunities for RPA binding to the G-strand: (1) discontinuous lagging strand synthesis during DNA replication through telomeric regions, (2) transcription of TERRA RNA (reviewed in^41^), and (3) the ssDNA overhang at the end of the telomere (**Figure 7**). While the two former events are transient, important during early S-phase and depend on the RPA ability to melt the telomeric G4 structures, the latter is the site of competition between RPA and POT1.

**Figure 7.**
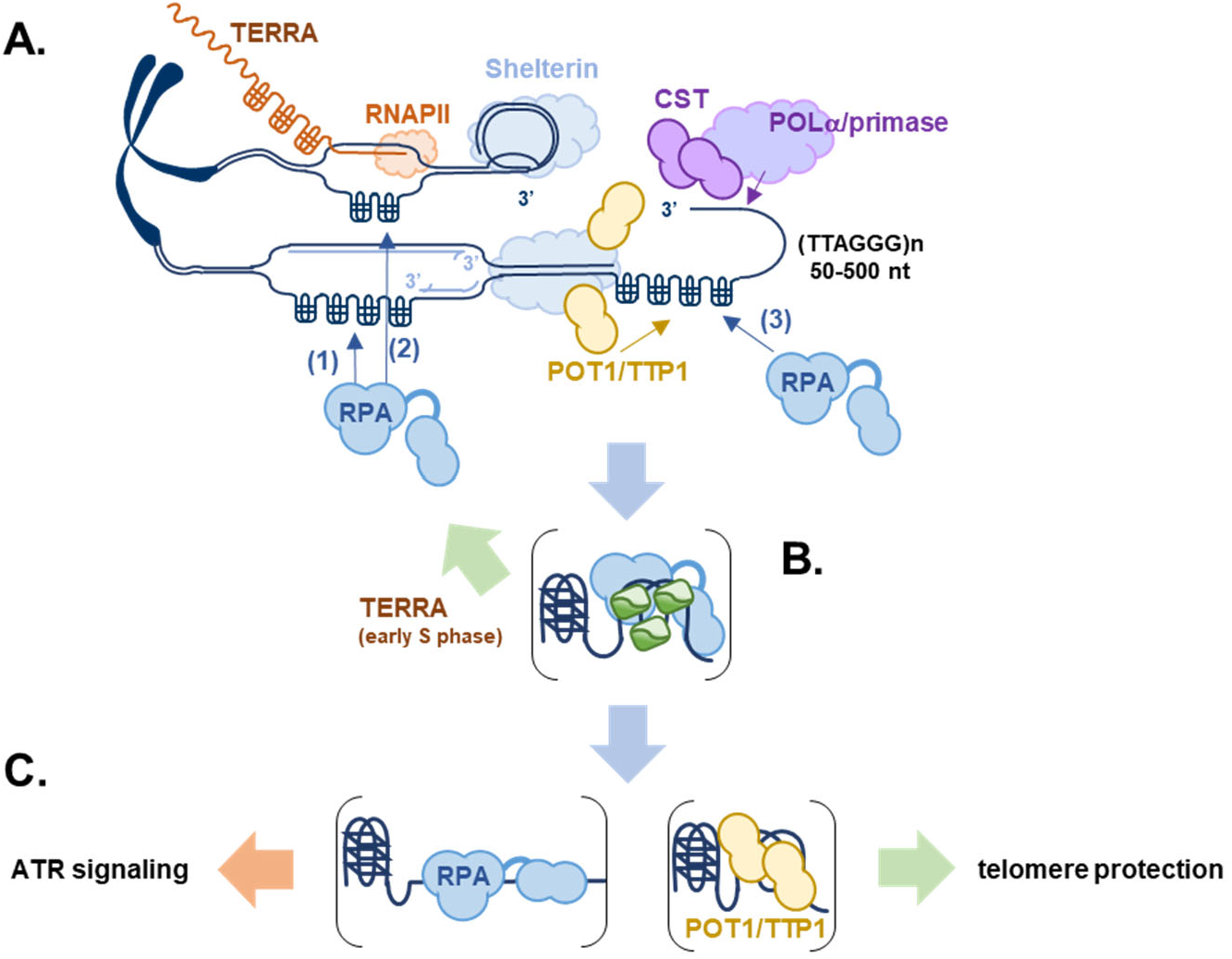
Telomere-specific RPA-DNA-hnRNPA1 complex. **A.** Several distinct events at human telomeres may lead to formation of G-quadruplexes that require RPA presence: (1) replication of the telomeric DNA and (2) transcription of TERRA RNA expose the G-strand, while (3) the end of human telomeres is a 50-500 nt 3′ ssDNA overhang. **B**. Formation of the RPA-DNA-hnRNPA1 complex is specific to telomeric G-rich DNA and may play a protective role until RPA is replaced at the ssDNA overhangs with telomere-specific POT1/TTP1. Stable RPA-hnRNAPA1 complex with constrained conformational dynamics may present a nucleoprotein structure refractory to DNA damage signaling (**C**).

RPA retained at telomeric ssDNA may elicit a DNA damage response via ATR kinase, which responds to DNA lesions that have been processed into ssDNA-RPA intermediates^42–44^ and plays an important role in telomere maintenance^45,46^. Both POT1 and RPA have high affinity for telomeric ssDNA, but POT1 is over 100-fold less abundant in cells compared to RPA^47^. RPA-mediated G-quadruplex melting is also important for binding of another telomere-specific ssDNA binding complex CTC1-STN1-TEN1 (CST) whose binding to telomeric ssDNA is inhibited by the formation of G-quadruplex structures^48^. Recent structural and functional analyses showed that CST is recruited to the telomeric ssDNA by POT1/TTP1, and in turn recruits POLα/primase complex to initiate fill-in synthesis of the C-strand^49^. One or several of these finely tuned nucleoprotein transactions involving RPA may be affected by the tighter telomeric ssDNA engagement and altered dynamics of the DBD-A in RPA^E240K^ resulting in the telomere specific defect.

The mechanisms by which RPA targets replicating DNA and the displaced strand of the transcription bubble, while POT1 protects the telomeric ssDNA overhang against RPA-mediated ATR signaling remains unclear. Here, we propose that the exchange of RPA for POT1-TPP1 occurs via a ternary complex where telomeric ssDNA simultaneously is occupied by RPA and hnRNPA1. Previously it was suggested that hnRNPA1 competes with RPA^21^. Instead, we observe formation of a ternary complex, where both RPA and hnRNPA1 are bound to the same telomeric sequence. While the complex is stable persisting for many minutes, RPA-DNA contacts are rearranged in the ternary complex with both DBD-A and DBD-D being only partially engaged. Modular organization of the RPA DBDs allows for the flexibility of the telomeric ssDNA containing both RPA and hnRNPA1, and from the DNA perspective, the ternary complex is more dynamic than h-telG4-RPA complex. This remodeling of the RPA-DNA contacts parallels that observed for yeast RPA and recombination mediator Rad52, which upon binding to RPA-ssDNA complex modulates the contacts between RPA DBD-D and ssDNA resulting in a stable RPA-DNA-Rad52 complex which nevertheless provides binding platform for Rad51 recombinase without exposing ssDNA^3^.

RPA was recently shown to form condensates on ssDNA that facilitate telomere maintenance^50^. Another study, however, showed that RPA is excluded from telomeric condensates while POT1 is recruited^51^. In addition to its RNA-binding domains, hnRNPA1 contains a low complexity sequence domain at the C-terminus that endows hnRNPA1 with a propensity to undergo liquid-liquid phase separation^52^. It is tempting to speculate that both, phase separation and/or configuration of the ternary RPA-DNA-hnRNPA1 complex may restrict ATR access to the RPA -coated telomeric DNA and subsequent DNA damage signaling (**Figure 7**).

While unexpected, the ability of TERRA RNA to more readily strip hnRNPA1 from the RPA-telomeric ssDNA-hnRNPA1 ternary complex as compared to the binary hnRNPA1-telomeric ssDNA complex makes physiological sense. A broad specificity of hnRNPA1 for different RNA sequences^53,54^ underlies its many functions in cellular RNA metabolism which include regulation of alternative RNA splicing, mRNA transcription and translation, and RNA stability (reviewed in ^55^). The preferred hnRNPA1 binding sequence identified by SELEX, UAGGG(A/U) resembles telomeric ssDNA, TERRA RNA, and consensus sequences of vertebrate splice sites^40^. Structurally, hnRNPA1 binding to and bridging RNA segments is likely to involve a similar set of contacts as its complex with telomeric ssDNA^30^. If TERRA were to disrupt all hnRNPA1-RNA/DNA contacts, it would have a deleterious effect on RNA metabolism. It is not surprising therefore, that the contacts between hnRNPA1 and telomeric ssDNA are different in the presence and absence of RPA.

It is notable that while RPA and hnRNPA1 physically interact^37^, the ternary RPA-DNA-hnRNPA1 complex is specific to telomeric G-quadruplex DNA, as hnRNPA1 was unable to remodel RPA bound to non-structured ssDNA, or to G-quadruplexes formed in the promoters of *MYC* and *BCL-2* genes. These non-telomeric quadruplexes play important regulatory functions in gene expression and can also have pathological roles by interfering with DNA replication and repair. RPA presence at and melting of these G-quadruplexes is likely controlled by a range of G-quadruplex binding and unfolding factors. Formation of a stable RPA-hnRNPA1 complex would interfere with the function of these DNA structures. Similarly, hnRNPA1 removal from the ternary complex containing RPA and telomeric ssDNA by numerous hnRNPA1-binding cellular or viral RNAs would drastically reduce the chance of formation of the telomere specific RPA-DNA-hnRNPA1 complex.

## Methods

### Chemicals and reagents

All chemicals were reagent grade (Sigma-Aldrich, St. Louis, MO). All fluorophores used to generate fluorescently labeled proteins were purchased from Click Chemistry Tools. Cy3-labeled, Cy-5 labeled, biotinylated and unmodified oligonucleotides were purchased from Integrated DNA Technologies. Sequences of all DNA oligonucleotides are listed in the **Supplementary Table 1**. Twelve mM Trolox (6-hydroxy-2,5,7,8-tetramethylchromane-2-carboxylic acid, Sigma-Aldrich; 238813-1G) solution was prepared as described previously^3^ by adding 60 mg of Trolox (238813-5G, Sigma-Aldrich) to 10 mL of water with 60 μL of 2 M NaOH, mixing for 3 days, filtering, and storing at 4 °C. Oxygen scavenging system, Gloxy was prepared as a mixture of 4 mg/mL catalase (C40-500MG, Sigma-Aldrich) and 100 mg/mL glucose oxidase (G2133-50KU, Sigma-Aldrich) in K100, T100 or L100 buffer (see below).

### Protein Expression, labeling and generation of fluorescently labeled RPA variants

Both wild type RPA and mutant proteins were expressed and purified as previously described^9,56^. A pET28a(+) plasmid containing an open reading frame for human hnRNPA1 with an N-terminal 6xHis tag was synthesized by GenScript. The 6xHis-hnRNPA1 was expressed in *E. coli* Rosetta cells using IPTG (0.5 mM) induction for 3 h at 37 °C. The cells were then harvested and lysed by sonication in lysis buffer (400 mM NaCl, 20mM imidazole, 1.72M sucrose, 50mM KPi buffer pH 7.4, 5mM BME, 0.5% Triton X-100, and 0.05% w/v lysozyme). The protein was purified using metal-affinity chromatography (HisTrap HP, Cytivia). Fractions containing hnRNPA1 were loaded onto a Heparin Sepharose column (HiTrap HP, Cytivia) and eluted with a 30 mL gradient of 100 mM to 1 M NaCl in Heparin Buffer C containing 25 mM HEPES (pH 7.8), 0.1 mM EDTA, 1 mM DTT and 5% Glycerol. Peak fractions were then dialyzed against Heparin Buffer C containing 100 mM NaCl. RPA (wild type and mutant) and hnRNPA1 protein concentrations were determined by measuring absorbance at 280 nm using extinction coefficients of 88,830 M^-1^ cm^-1^ and 23,380 ^-1^ cm^-1^, respectively.

4AZP-incorporated RPA proteins were purified and labeled with MB543 as previously described^24^. For human RPA, 4AZP was positioned at either Ser-215 (DBD-A; RPA1) or Trp-107 (DBD-D; RPA2), respectively. Briefly, ∼3 ml of RPA4-AZP (10 μM) was incubated on a rocker with a 1.5-fold molar excess (15 μM) of dibenzocyclooctyne-amine fluorophore (DBCO-MB543, Click Chemistry Tools Inc.) for 2 h at 4 °C. Labeled RPA variants were separated from excess dye using a Biogel-P4 gel filtration column (Bio-Rad Laboratories; 65 ml bed volume) using storage buffer (30 mM HEPES, pH 7.8, 200 mM KCl and 10% (v/v) glycerol). Fractions containing labeled RPA were pooled, concentrated using a 30-kDa cut-off spin concentrator, and flash-frozen using liquid nitrogen. Fluorescent RPA was stored at −80 °C. Labeling efficiency was calculated using the respective extinction coefficients (ε) ε280 = 87,410 M^−1^ cm^−1^ for RPA and ε550 = 105,000 M^−1^ cm^−1^ for DBCO-MB543. We obtained 45 ± 17% and 40 ± 25% labeling efficiencies for the RPA–DBD-A^MB543^ and RPA–DBD-D^MB543^, respectively.

### Single-Molecule TIRFM

All single-molecule TIRFM studies were performed using a prism-based microscope custom-built around an Olympus IX71 microscope frame^3,57^. Videos were recorded using an electron-multiplying charge-coupled device camera (Andor; DU-897-E-CSO-#BV) at 100-ms time resolution. Background was set to 400, correction was set to 1,200 and gain was set to 250 for all videos recorded. Quartz slides (25 mm × 75 mm × 1 mm #1x3x1MM, G. Frinkenbeiner, Inc.) and cover glass (24 mm × 60 mm-1.5, Fisherbrand) were washed, coated and flow cells were assembled as described previously^3,58^. Assembled flow cells were mounted onto the microscope stage and rinsed with T100 buffer (10 mM Tris-HCl, pH 7.5, and 50 mM NaCl), K100 buffer (10 mM Tris-HCl, pH 7.5, and 100 mM KCl), or L100 buffer (10 mM Tris-HCl, pH 7.5, and 100 mM LiCl), then incubated with 0.2 mg ml^−1^ NeutrAvidin (Thermo Fisher) for 3 min and rinsed with buffer again.

Prior to surface tethering, the mixture biotinylated DNA oligo and human telomeric G4-forming oligo (see **Supplementary Table 1**) were heated together at 95 °C for 5 min and slowly cooled down to allow for annealing and G4 folding, and were later diluted to indicated working concentrations in the imaging buffers described below. Before starting each experiment, the DNA was refolded and reannealed to ensure proper folding.

DNA tethering and imaging was carried out in the following buffers: Poly-dT ssDNA binding experiments were carried out in reaction buffer containing 50 mM Tris-HCl, pH 7.5, 5 mM MgCl_2_, 100 mM NaCl, 1 mM DTT, 1 mg/ml BSA, 0.8% w/v D-glucose, 12 µM glucose oxidase, 0.04 mg/ml catalase in Trolox solution. Telomeric G-quadruplex binding and smFRET experiments were carried out in reaction buffer containing 50 mM Tris-HCl, pH 7.5, 5 mM MgCl_2_, 100 mM KCl or LiCl, 1 mM DTT, 1 mg/ml BSA and 0.8% w/v D-glucose, 12 µM glucose oxidase, 0.04 mg/ml catalase in Trolox solution. To tether DNA to the slide surface, the flow cell was incubated for 3 min with 100 pM of biotinylated d100T or 1pM of biotinylated G-quadruplex (hTelG4-comp annealed to biotin base and pre-folded in either KCl or LiCl-containing buffer; see **Supplementary Table 1**) in the respective reaction buffer, then rinsed with reaction buffer to remove untethered DNA. Movies were recorded for 210 seconds at a frame rate of 100 ms. Wild type RPA or RPA^E240K^ labeled with MB543 in DBD-A or DBD-D was added at an indicated concentration after the first 30 seconds. At 120 seconds, free RPA was removed with reaction buffer or replaced with 50 nM hnRNPA1.

The effect of TERRA RNA was monitored by collecting 100 frame movies at a frame rate of 100ms after each substrate addition (100 pM DNA, 100 pM RPA, 50 nM hnRNPA1, and 50 nM TERRA RNA) within 5 different locations on the slide surface. Representative frames depicting portions of fields of view were color-inverted in ImageJ and cropped to 1/4 of the field of view. The number of trajectories with a fluorescence signal above background were quantified for 5 movies collected under the same condition. The fluorescence intensity of each point in these trajectories was calculated and plotted. Statistical analysis was performed using ordinary one-way ANOVA within GraphPad Prism.

Single-molecule FRET data were recorded in two independent experiments for both KCl and LiCl conditions. First the DNA alone was recorded followed by the addition of different concentration of indicated proteins. In the presence of KCl, proteins, once flowed in cell, were incubated for 8 mins and for LiCl proteins were incubated for 2 mins. FRET efficiency histograms were obtained by in-house MATLAB program using 10 different slide location data sets (each location contains 200–600 single molecules) for each condition. FRET was calculated as the ratio between the acceptor intensity and the sum of the acceptor and donor intensities after donor leakage correction to Cy5 channel^59–61^. Calculated FRET values for each molecule in the selected movies were binned with the bin size of 0.02 and plotted using GraphPad Prism.

### Single-Molecule analysis of RPA conformational dynamics

An IDL script was used to extract fluorescence intensity trajectories from each video as previously described^3^. Trajectories were viewed and selected for analysis using a Matlab script. Only those trajectories that showed the appearance of the fluorescence signal between 30 s and 120 s, and not during the first 30 s, and had a signal-to-noise ratio > 4 (raw signal) were selected for analysis. The selected trajectories were then saved individually and globally processed for analysis by hFRET^26^. Idealized trajectories were imported from hFRET to KERA Matlab script^62^, which was used to optimize state assignment to avoid overfitting and remove transient events (1 or 2 frames in duration). Dwell times for each state were binned, plotted as frequency distributions and fitted to exponential functions using GraphPad Prism. Transition Density Plots (TDPs) were generated from the idealized fluorescence trajectories using a custom Matlab script, which extracts the raw idealized states and translons from the hFRET data. The resulting 3D histogram is then overplayed with a Gaussian smoothing algorithm to minimize variation and emphasize trends within the hFRET transitions^63^.

### Bulk FRET measurements

After the baseline of buffer only was recorded (30 mM Tris-HCl pH 7.5, 100 mM LiCl and 1 mM DTT or 30 mM Tris-HCl pH 7.5, 1 mM DTT, 5 mM MgOAc_2_ and 100 mM KCl), 10 nM of dT15, h-tel-15, dT30, h-telG4, BCL-2G4 1245, or PU27G4 (**Supplementary Table 1)** was added to the cuvette followed by addition of indicated concentrations of the wild type or mutant RPA, hnRNPA1, and/or TERRA RNA, SELEX RNA, or HIV ESS3 RNA (**Supplementary Table 1**).

FRET efficiency was calculated as 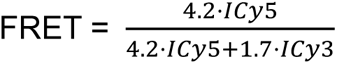, where *ICy5* is the averaged acceptor intensity and *Icy3* is the averaged donor intensity after subtracting the background fluorescence. The calculated FRET efficiency was plotted against protein or RNA concentrations and analyzed in GraphPad Prism. Equilibrium binding curves were fitted using quadratic binding equation to determine the K_d_s.

***Mass photometry*** of the RPA (100 nM), hnRNPA1 (600 nM), and their complexes with each other and telomeric DNA (100 nM) was performed using the Refeyn TwoMP mass photometry instrument (Refeyn Ltd. Oxford, UK) in buffer containing 20mM Tris (pH 7.4), 100mM KCl, and 1 mM DTT. Cover slides and silicon buffer gaskets were washed twice with miliQ water, 100% isopropanol, and again with miliQ water, and dried under an air stream at room temperature. Dried silicon gaskets were attached to the glass slide by applying a mild pressure and mounted. Molecular weight calibrations were performed using two protein oligomer solutions, β-amylase (56, 112 and 224 kDa) and Thyroglobulin (670 kDa). In each experiment, the indicated proteins and DNA were mixed to form 4x solution at room temperature, and then diluted 4-fold into the buffer-filled gasket. Individual molecular weights collected from 3000 frames (59.9 seconds) were binned in 3 kDa bins and plotted as frequency histograms. GraphPad Prism was used to fit the molecular weight distributions to multiple Gaussians.

### Electrophoretic Mobility Shift Assays (EMSAs)

DNA substrates (h-telG4, BCL-2 G4, or Pu27G4) were allowed to fold into G-quadruplexes by heating 1µM DNA solution at 95°C for 5 minutes followed by slow equilibration of the heating block to room temperature. Folded G-quadruplexes (30 nM molecules) or dT_30_ ssDNA were incubated with indicated amounts of RPA, hnRPA1 and RPA+hnRPA1 at 4°C for 30 min in 20 µl of standard reaction buffer, containing 20 mM Tris-Acetate (pH 7.5), 100 mM KCl, 1 mM EDTA, 5mM MgCl_2_, 10% glycerol, 0.1 mg/ml BSA and 5 mM DTT. Two µl of the loading dye solution (20 mM Tris-Acetate (pH 8.0), 1 mM EDTA, 10% glycerol and 0.25 % Orange-G (w/v)) was added for each reaction. The reaction products were separated by electrophoresis on the 4.8% (29:1) native polyacrylamide gel in 20 mM Tris-Acetate (pH 8.0), 1 mM EDTA and 50mM KCl buffer. The gels were visualized separately in Cy3 and Cy5 channels using the ChemiDoc MP imaging system (BIO-RAD).

### Co-immunoprecipitation and immunoblot

Previously established induced pluripotent stem cells (iPSCs) cell lines with wild type RPA1 (RPA1^E240^) and RPA1 p.E240K (RPA1^E240K^)^9^ were lysed in RIPA lysis buffer (Cat# 20-188 ) supplemented with protease and phosphatase inhibitor cocktail (Thermo Cat# A32961 on ice for 20 minutes followed by centrifugation (14,000 × g, 10 min, 4 °C). The supernatant was removed and stored on ice while the remaining cell pellets were digested with micrococcal nuclease (BioLabs Cat no. M0247S) to extract chromatin bound nuclear complexes, and centrifuged (14,000 × g, 10 min, 37 °C). Supernatants from RIPA and micrococcal nuclease digestion were combined, pre cleared for 20 minutes with agarose IgG beads (Sigma Cat# A0919) and incubated with 1 μg of anti-hnRNPA1 antibody (Santa Cruz Biotech, sc-32301) overnight at 4 °C with constant rotation. Protein A/G agarose beads (Thermo, Cat no. 88802) were used for pulldown according to the manufacturer’s protocol. Precipitates were washed four times with cold lysis buffer, then resuspended in lysis buffer with SDS sample loading buffer, boiled for 10 min, and immediately subjected to SDS-PAGE for immunoblotting. Images were obtained using ChemiDoc Imaging System (BioRad). Anti-FLAG (Sigma, F1804, 1:1000) antibody was used for IgG control. Membranes were immunoblotted with anti-RPA1 (ThermoFisher, MA5-36226, 1:1000), anti-hnRNPA1 antibody (Santa Cruz Biotech, sc-32301) and anti-H3 (Cell Signaling, Cat no. 9717, 1:1000) antibodies.

## Author Contributions

SLG: smTIRFM and FRET-based experiments and data analyses, conceptual design of the study; RS: cell-based studies, conceptual design of the study; VK: preparation and characterization of fluorescently-labeled RPA variants; MR: mass photometry; MH: EMSAs, smTIRFM data analysis; PG: smFRET; DB: bulk FRET-based experiments; SML, JEM, BAW and SMAT: single-molecule data analysis; MW: conceptual design; EA: fluorescently-labeled RPA variants, data interpretation; MS: conceptual design, data analysis and interpretation. All authors contributed to manuscript preparation and editing.

## Data and Materials Availability

Data, plasmids for protein expression, and code for single-molecule data analysis are available from the corresponding author upon request.

