## Supplemental Information for "Human hnRNPA1 reorganizes telomere-bound Replication Protein A"

### Supplementary Table 1. Oligonucleotides Used in This Study

| Name | Sequence | Experiment |
|---|---|---|
| d100T | 5'-bio-(T)$_{100}$-3' | smTIRFM with MB543-labeled RPAs |
| biotin-base | 5'- GCCTCGCTGCCGTCGCCA -bio-3' | smTIRFM, surface-tethering |
| biotin-base (Cy5) | 5'-**Cy5**-GCCTCGCTGCCGTCGCCA -bio-3' | smTIRFM, surface-tethering for smFRET |
| h-telG4-comp | 5'-TGGCGACGGCAGCGAGGC-TTA-(<u>GGG</u>TTA)$_4$-3' | smTIRFM, five telometric repeats with 18 nt complementary to the biotin-base |
| h-telG4-comp (Cy3) | 5'-TGGCGACGGCAGCGAGGC-TTA-(<u>GGG</u>TTA)$_4$-**Cy3**-3' | smTIRFM (smFRET), five telometric repeats with 18 nt complementary to the biotin-base |
| h-tel-15(FRET) | 5'-**Cy5**-TTAGGGTTAGGGTTA-**Cy3**-3' | Bulk FRET |
| dT30 | 5'-**Cy5**-(T)$_{30}$**Cy3**-3' | Bulk FRET; EMSA |
| h-telG4(FRET) | 5'-**Cy5**-(TTA<u>GGG</u>)$_5$-**Cy3**-3' | Bulk FRET; EMSA |
| h-telG4 | 5'-(TTA<u>GGG</u>)$_5$-3' | MP |
| BCL-2G4 | 5'-A<u>GGGG</u>C<u>GGG</u>CGCTTTAGGAAA<u>GGG</u>C<u>GGG</u>T-3' | Bulk FRET |
| PU27G4 | 5'- T<u>GGGG</u>A<u>GGG</u>T<u>GGGG</u>A<u>GGG</u>T<u>GGGG</u>AAGG-3' | Bulk FRET |
| TERRA RNA | 5'-(UUAGGG)$_3$-3 | Bulk FRET |
| SELEX RNA | 5'-UAUGAUAGGGACUUAGGGUUAGGAU-3' | Bulk FRET |
| HIV ESS3 RNA | 5'-GAUCGAUUCGAUUAGUGAGCGGAUU-3' | Bulk FRET |

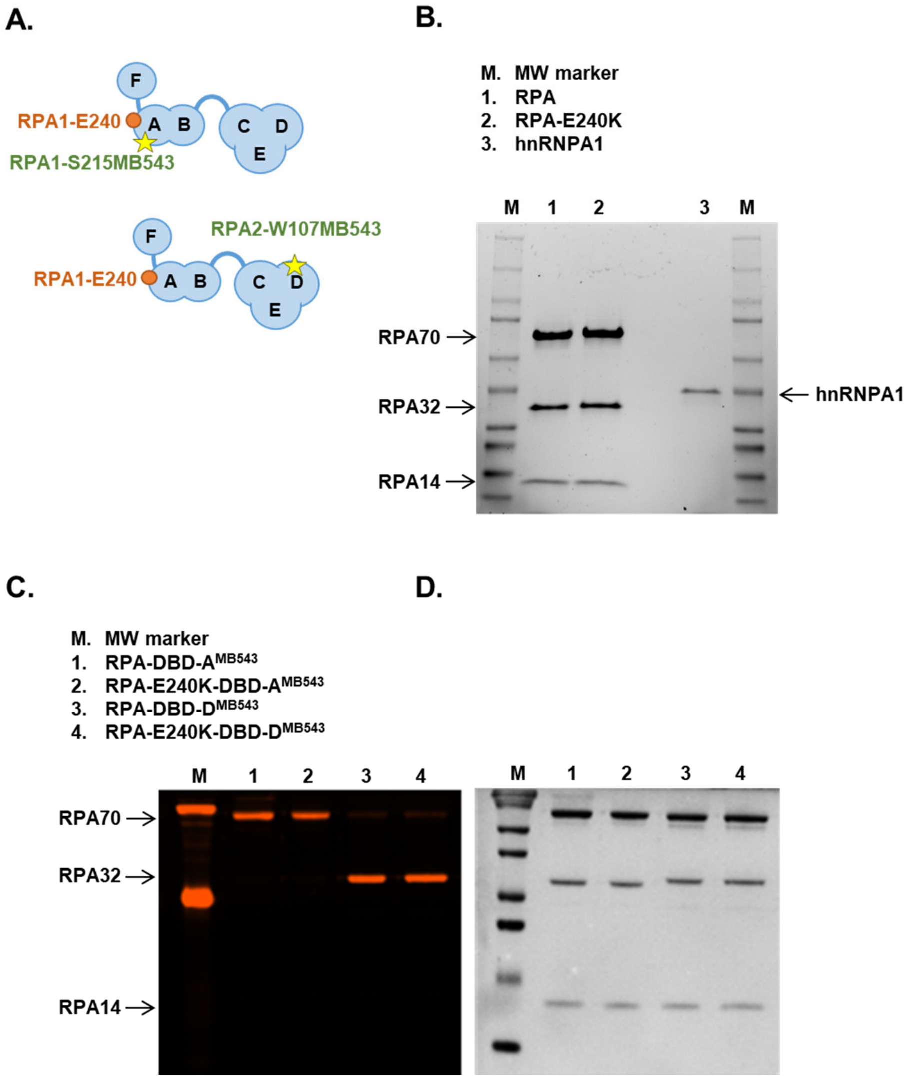


**Supplementary Figure 1. Recombinant proteins used in this study. A.** Schematic representation of human RPA protein with locations of the E240K mutation in RPA1 (RPA70) and of the MB543 label in the DBD-A of RPA1 (RPA70) or DBD-D of RPA2 (RPA32). **B.** CBB-stained SDS-PAGE gel of unlabeled purified proteins. **C**. Fluorescent image of MB543-labeled RPAs resolved by PAGE. **D**. The same gel stained with CBB.

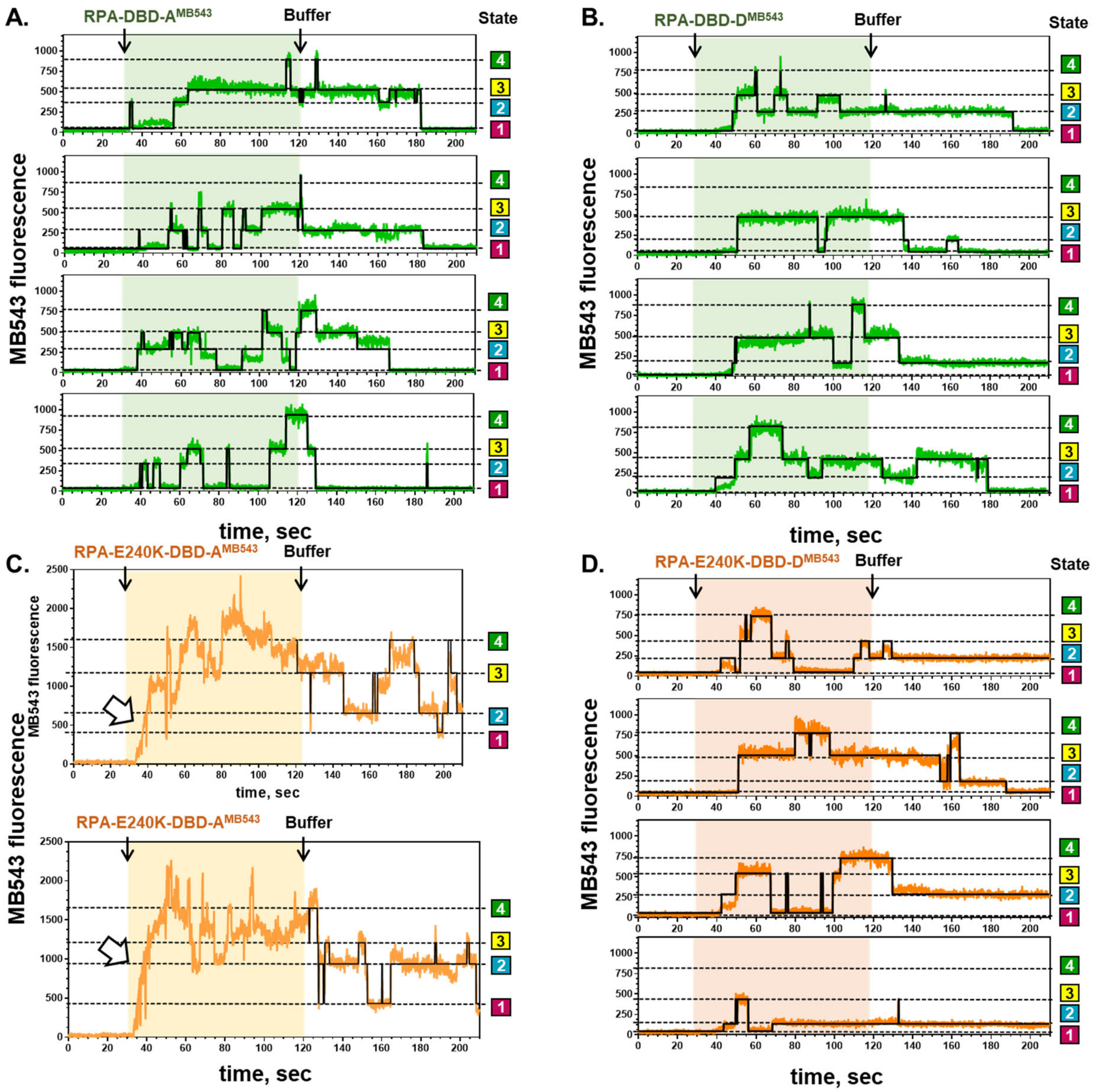


**Supplementary Figure 2** (Related to Figure 1)**. Additional representative fluorescence trajectories** for the RPA-DBD-$A^{MB543}$(**A**), RPA-DBD-$D^{MB543}$ (green) (**B**), $RPA^{E240K}$-DBD-$A^{MB543}$ (**C**), $RPA^{E240K}$-DBD-$D^{MB543}$ (orange) (**D**) binding to unstructured ssDNA overlaid with idealized trajectories (black) obtained by globally fitting all trajectories to a four-state model using hFRET.

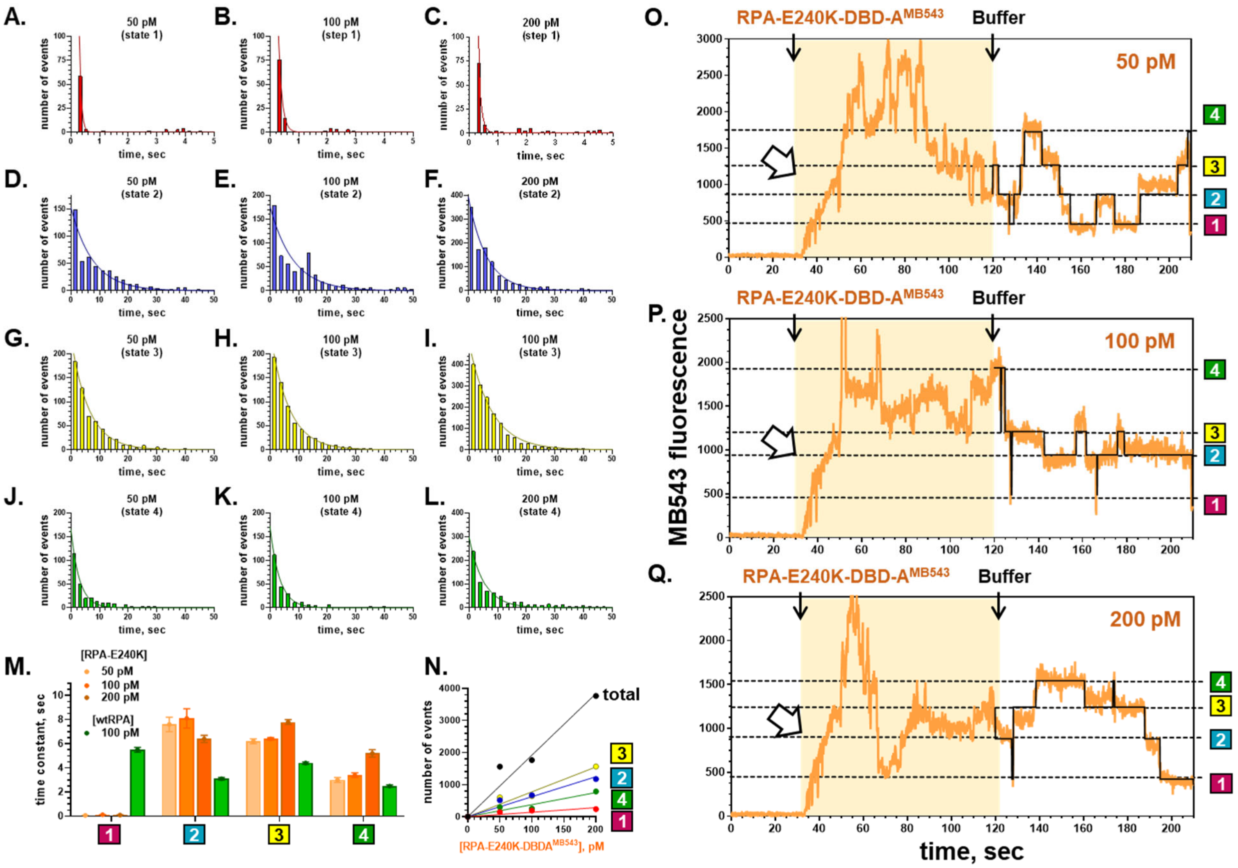


**Supplementary Figure 3** (Related to Figure 1)**. Individual molecules of RPA$^{E240K}$-DBD-A$^{MB543}$ exhibit an idiosyncratic fluorescence pattern upon binding to unstructured ssDNA. A-L.** Dwell time analysis of smTIRFM experiments carried out with 50, 100 and 200 pM RPA$^{E240K}$-DBD-A$^{MB543}$ in the reaction chamber. The dwell times for states 2, 3 and 4 were binned with a bin size of 2.4 seconds (blue, yellow and green bars, respectively), while dwell times for state 1 were binned with a bin size 0.2 second (red bars). Solid lines represent exponential fits for each distribution. **M.** Histogram plot of the time constants from the exponential fitting of the dwell time distributions ± fitting error. Orange bars show time constants for different concentrations of the RPA$^{E240K}$-DBD-A$^{MB543}$, green bars show time constants derived from the dwell time distributions for RPA$^{E240K}$-DBD-D$^{MB543}$. Note the absence of any detectable trend in concentration dependence of the RPA$^{E240K}$-DBD-A$^{MB543}$ dwell times. **N**. Number of fluorescent trajectories extracted from the RPA$^{E240K}$-DBD-A$^{MB543}$ and the number of identified states increased linearly with increasing of the protein concentration. The data shown in **M** and **N** confirm that fluorescence trajectories originate from individual RPA$^{E240K}$-DBD-A$^{MB543}$ molecules macroscopically bound to ssDNA and that the idiosyncratic increase in the MB543 fluorescence upon initial binding stems from the binding mode of this RPA mutant.

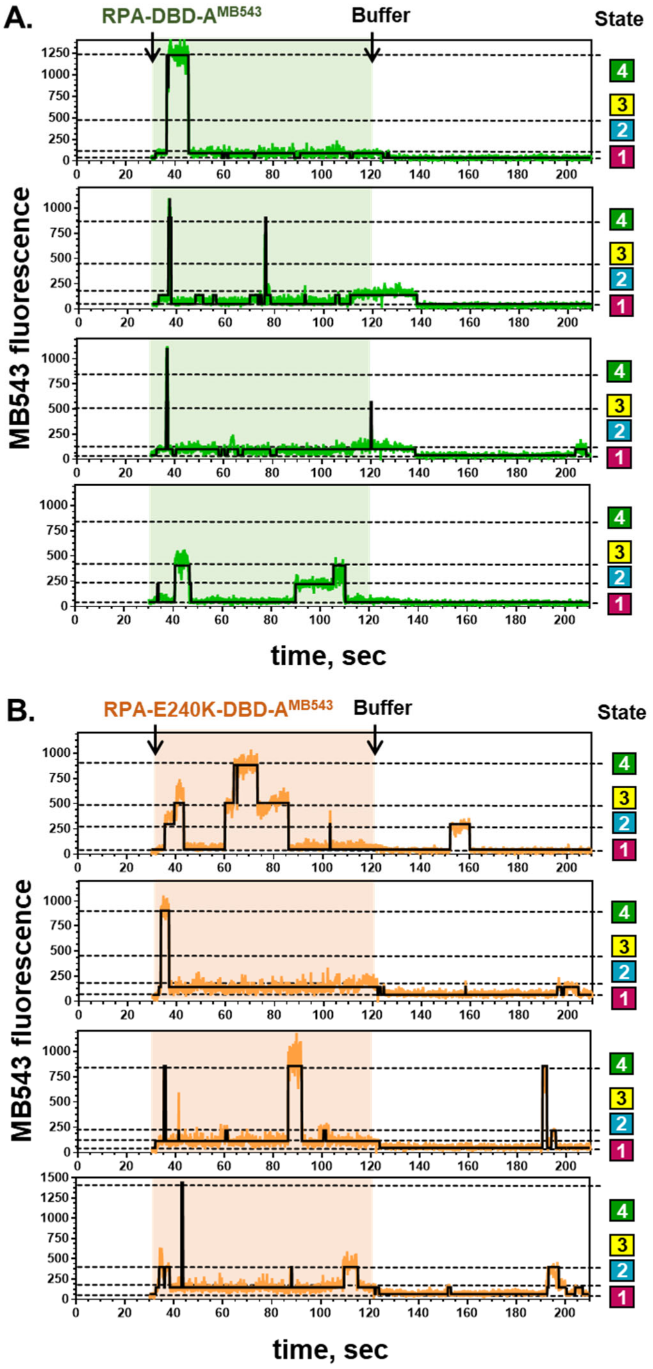


**Supplementary Figure 4** (Related to Figure 2)**. Additional representative fluorescence trajectories** for the RPA-DBD-A$^{MB543}$(**A**) (green) and RPA$^{E240K}$-DBD-A$^{MB543}$ (**B**) (orange) binding to the folded telomeric G-quadruple  overlaid with idealized trajectories (black) obtained by globally fitting all trajectories to a four-state model using hFRET.

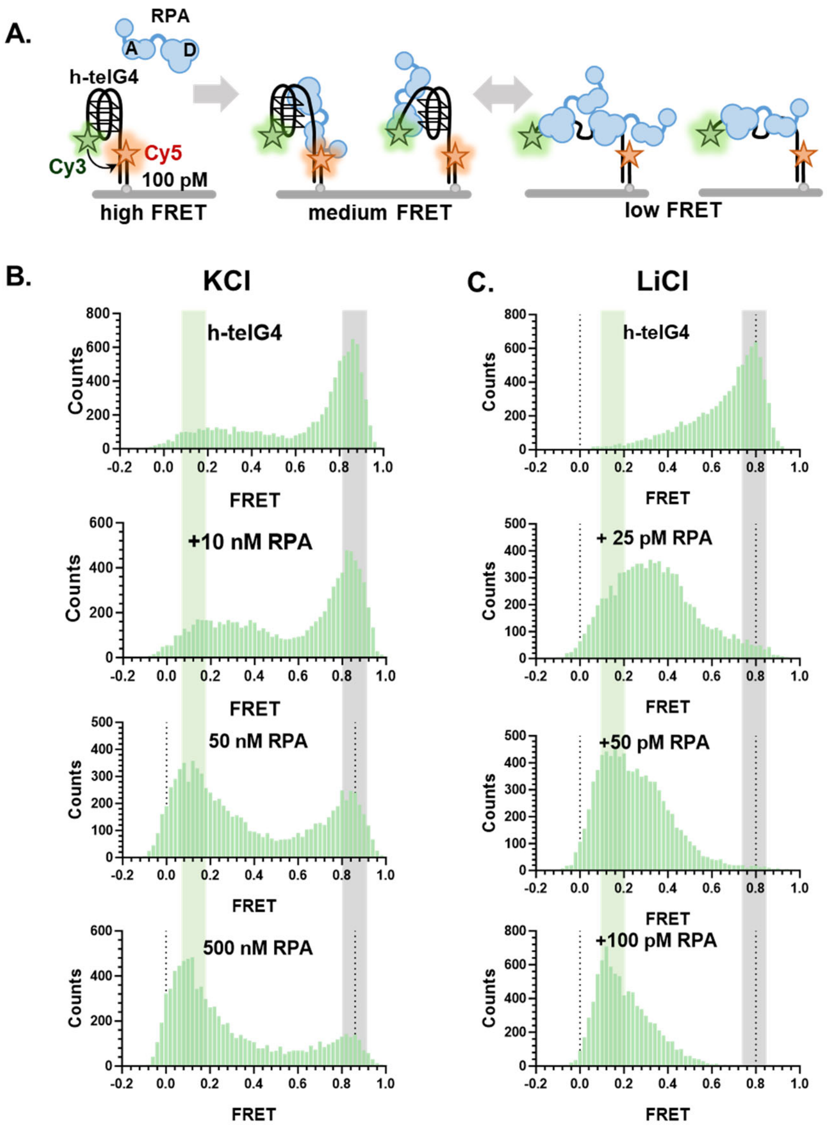


**Supplementary Figure 5** (Related to Figure 2)**. Telomeric G-quadruplex unfolding by RPA in the presence of potassium and lithium. A.** Experimental scheme for the single-molecule FRET (smFRET) based analysis of the RPA binding to the surface-tethered h-telG4 DNA. Hundred pM of Cy3/Cy5-labeled partial duplex DNA with the h-telG4 pre-formed in $K^+$ or $Li^+$-containing buffer were tethered to the smTIRFM flow cell surface. Equilibrium distributions of the FRET states were obtained at the indicated concentrations of RPA in $K^+$- (**B**) or $Li^+$-containing (**C**) buffer. For all measurements, FRET values collected from 10 randomly chosen fields of view, binned in 0.02 unit bins and plotted as histograms. Experiments were repeated at least twice. A representative set of distributions is shown.

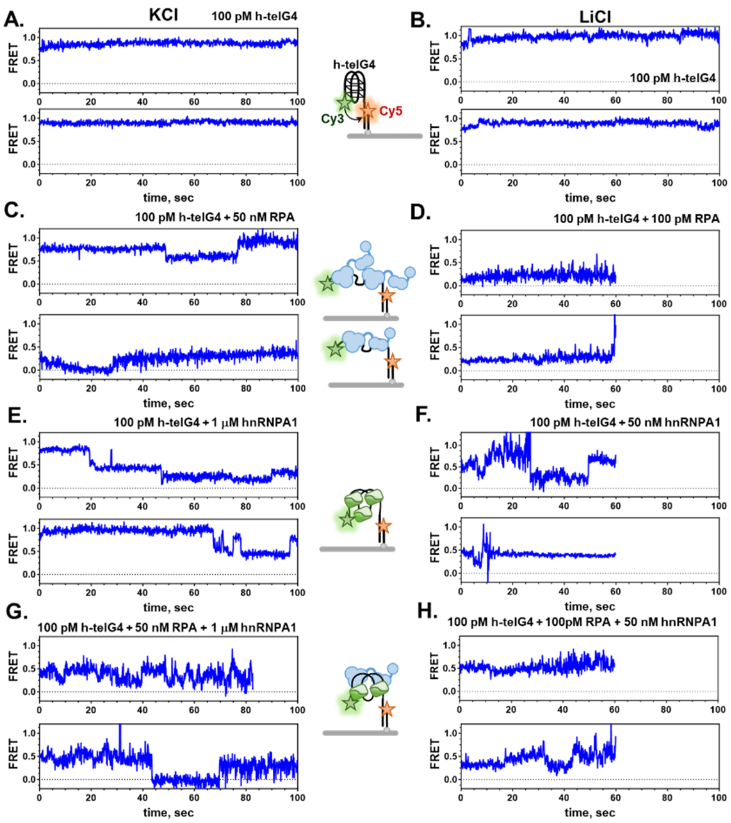


**Supplementary Figure 6** (Related to Figures 2 and 3)**. Representative smFRET trajectories** for the protein-free h-telG4 DNA in KCl (**A**) and LiCl (**B**); in the presence of saturating amounts of RPA (**C&D**), in the presence of hnRNPA1 (**E&F**); and in the presence of both RPA and hnRNPA1 (**G&H**).

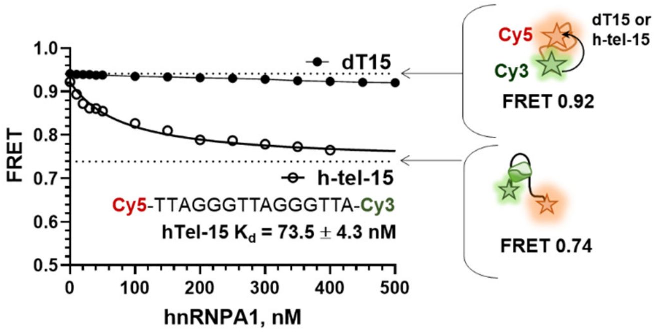


**Supplementary Figure 7** (Related to Figure 3)**. Purified hnRNPA1 preferentially binds to telomeric ssDNA.** Bulk FRET-based analysis of the hnRNPA1 binding to 10 nM 15-mer poly(dT) (filled circles) and telomeric DNA (open circles) labeled with Cy3 (FRET donor) and Cy5 (FRET acceptor) at the two termini. The free and bound DNA substrates are schematically shown on the right with their respective FRET values. The experiments were carried out in triplicates and the values are plotted as average ± standard deviation for the three independent titrations. Note that the error bars for most measurements were smaller than the symbols' sizes. The binding curve to hnRNPA1 and telomeric DNA was fitted to a quadratic binding equation. The calculated $K_d$ is shown with its fitting error.

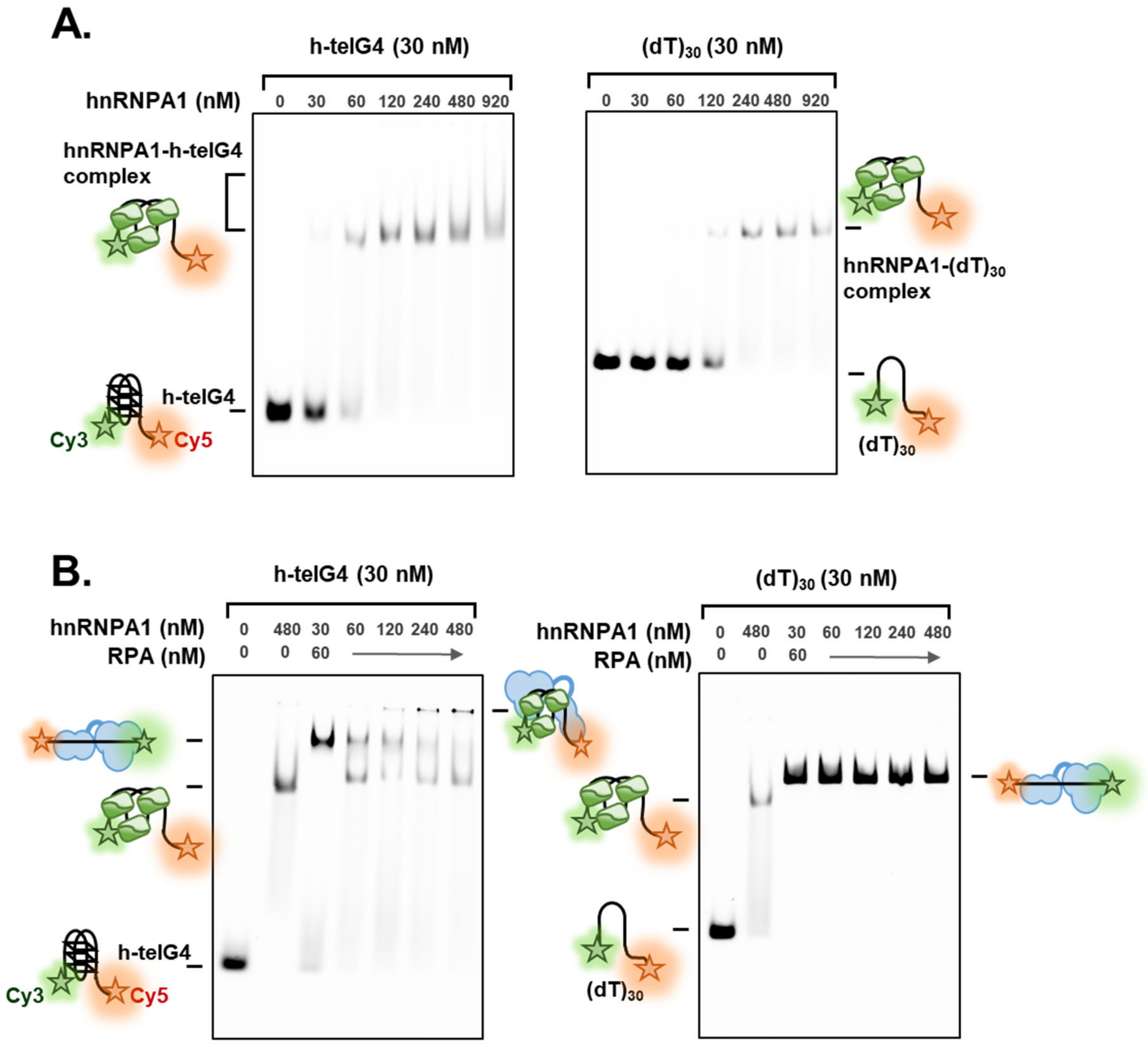


**Supplementary Figure 8** (Related to Figures 3 and 4)**. RPA and hnRNPA1 form ternary complex on telomeric ssDNA. A.** EMSA experiments showing change in the migration of the fluorescently-labeled DNA (30 nM) in the presence of indicated concentrations of hnRNPA1. While hnRNPA1 was able to bind unstructured ssDNA $(dT)_{30}$, much higher protein concentrations were needed to detect a complex compared to telomeric G-quadruplex. **B**. EMSA experiments showing appearance of the ternary RPA-DNA-hnRNPA1 complex when RPA was bound to telomeric G-quadruplex DNA. No super-shift was observed with unstructured DNA.

**Supplementary Table 2** (Related to Figure 4)**. Fitted molecular weights in the mass photometry experiments**

Calculated molecular weights:

RPA and $RPA^{E240K}$ ~ 111 kDa; hnRNPA1 ~ 38.7 kDa; h-telG4 ~ 8 kDa.

| panel | Experiment | Number of Gaussians | Measured molecular weights | |
|---|---|---|---|---|
| | | | Mean MW peak, kDa | Standard Deviation, kDa |
| **A.** | 50 nM RPA | 1 | 114.5 ± 0.1 | 11.1 ± 0.1 |
| **B.** | 50 nM $RPA^{E240K}$ | 1 | 118.6 ± 0.1 | 12.9 ± 0.1 |
| **C.** | 300 nM hnRNPA1 | 1 | 49.3 ± 0.1 | 10.9 ± 0.1 |
| **D.** | 50 nM RPA + 50 nM h-telG4 | 1 | 122.8 ± 0.1 | 10.6 ± 0.1 |
| **E.** | 50 nM $RPA^{E240K}$ + 50 nM h-telG4 | 1 | 126.7 ± 0.2 | 14.9 ± 0.2 |
| **F.** | 300 nM hnRNPA1 + 50 nM h-telG4 | 2 | 61.2 ± 0.2<br>86.0 ± 5.0 | 12.0 ± 0.4<br>22.1 ± 2.3 |
| **G.** | 50 nM RPA + 50 nM h-telG4 + 300 nM hnRNPA1 | 3 | 69.9 ± 1.0<br>124.3 ± 0.2<br>228.1 ± 1.2 | 15.1 ± 1.0<br>18.4 ± 0.2<br>33.3 ± 1.3 |
| **H.** | 50 nM $RPA^{E240K}$ + 50 nM h-telG4 + 300 nM hnRNPA1 | 3 | 73.1 ± 0.9<br>126.4 ± 0.1<br>209.2 ± 3.4 | 14.4 ± 0.9<br>18.7 ± 0.2<br>51.4 ± 3.2 |
| **I.** | 50 nM RPA + 300 nM hnRNPA1 | 3 | 59.7 ± 1.0<br>108.2 ± 0.1<br>139.2 ± 5.8 | 10.8 ± 1.4<br>17.4 ± 0.2<br>45.4 ± 5.8 |
| **J.** | 50 nM $RPA^{E240K}$ + 300 nM hnRNPA1 | 2 | 56.7 ±0.3<br>106.3 ±0.1 | 11.3 ± 0.4<br>16.0 ± 0.1 |

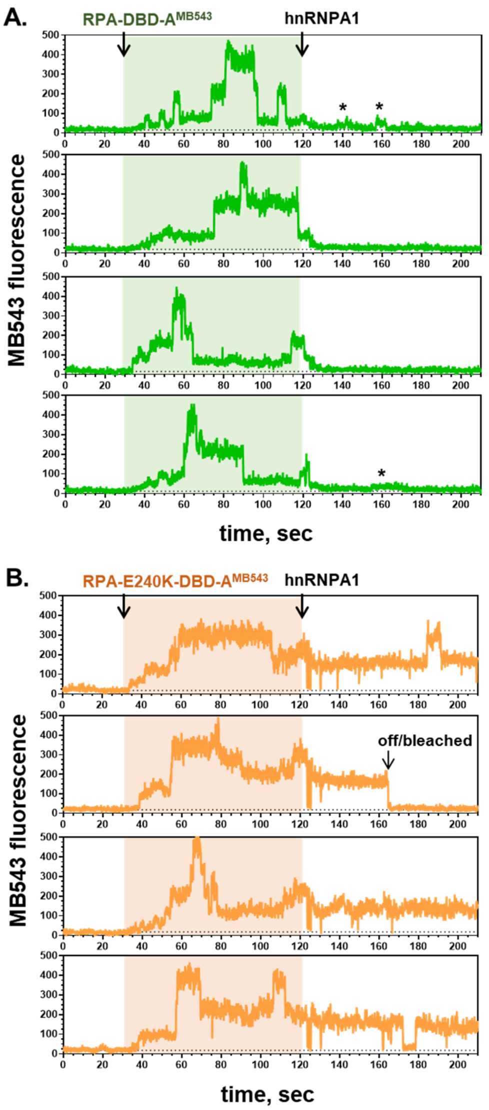


**Supplementary Figure 9** (Related to Figure 5)**. Additional representative fluorescence trajectories** for the RPA-DBD-A$^{MB543}$(**A**) (green) and RPA$^{E240K}$-DBD-A$^{MB543}$ (**B**) (orange) binding to the folded telomeric G-quadruple. At 30 seconds, the indicated fluorescently-labeled RPA was flown in, and at 120 seconds the unbound protein was replaced with 50 nM unlabeled hnRNPA1. Asterisks mark deviation of the fluorescence signal from the lowest state suggestive of the presence of RPA.

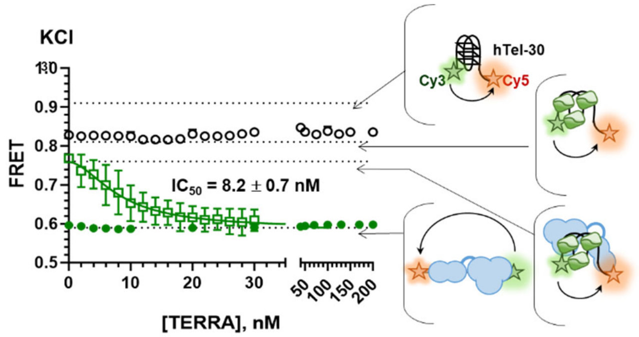


**Supplementary Figure 10** (Related to Figure 6)**. RPA-hnRNPA1 complex on telomeric ssDNA is remodeled by TERRA RNA.** Bulk FRET experiment. RPA-DNA (filled green circles), hnRNPA1-DNA (open black circles) and RPA-DNA-hnRNPA1 (open green squares) were preassembled on 10 nM of telomeric G-quadruplex folded in the presence of $K^+$ and challenged by addition of TERRA RNA at indicated concentrations. The experiments were carried out in triplicates and the values are plotted as average ± standard deviation for the three independent titrations. Note that the error bars for most measurements were smaller than the symbols' sizes. The curve for hnRNPA1 displacement from the ternary complex was fitted to an inhibition dose response equation. The apparent $IC_{50}$ value is shown with its fitting error.

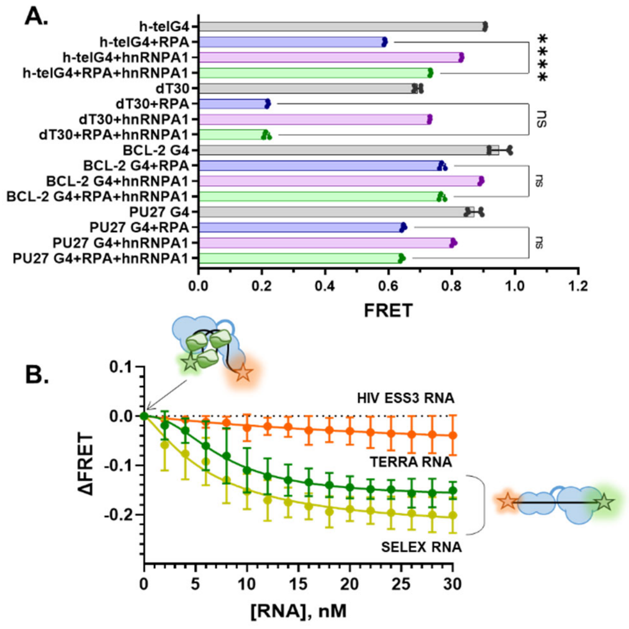


**Supplementary Figure S11. hnRNPA1-mediated RPA remodeling is specific to telomeric G-quadruplex and TERRA RNA. A.** Bulk FRET experiment of 10 nM RPA (blue bars), 200 nM hnRNPA1 (purple bars) and 10 nM RPA + 200 nM hnRNPA1 (green bars) binding to 10 nM of telomeric G-quadruplex folded in the presence of $K^+$ (hTelG4), dT30 ssDNA (in $Na^+$), BCL-2, and PU27 G-quadruplexes folded in the presence of $K^+$. Statistical analysis: ordinary one-way ANOVA; **** P<0.001, n.s. not significant. **B.** RPA-DNA-hnRNPA1 complex was preassembled on 10 nM of telomeric G-quadruplex folded in the presence of $K^+$ and challenged by addition of TERRA RNA (green circles), SELEX RNA (gold circles) or HIV ESS3 RNA (orange circles) at indicated concentrations (see Supplementary Table 1 for sequences of DNA and RNA oligos).

The data are plotted as change in FRET relative to the RPA-DNA-hnRNPA1 complex. The experiments were carried out at least in triplicates and the values are plotted as average ± standard deviation for the three or more independent titrations.

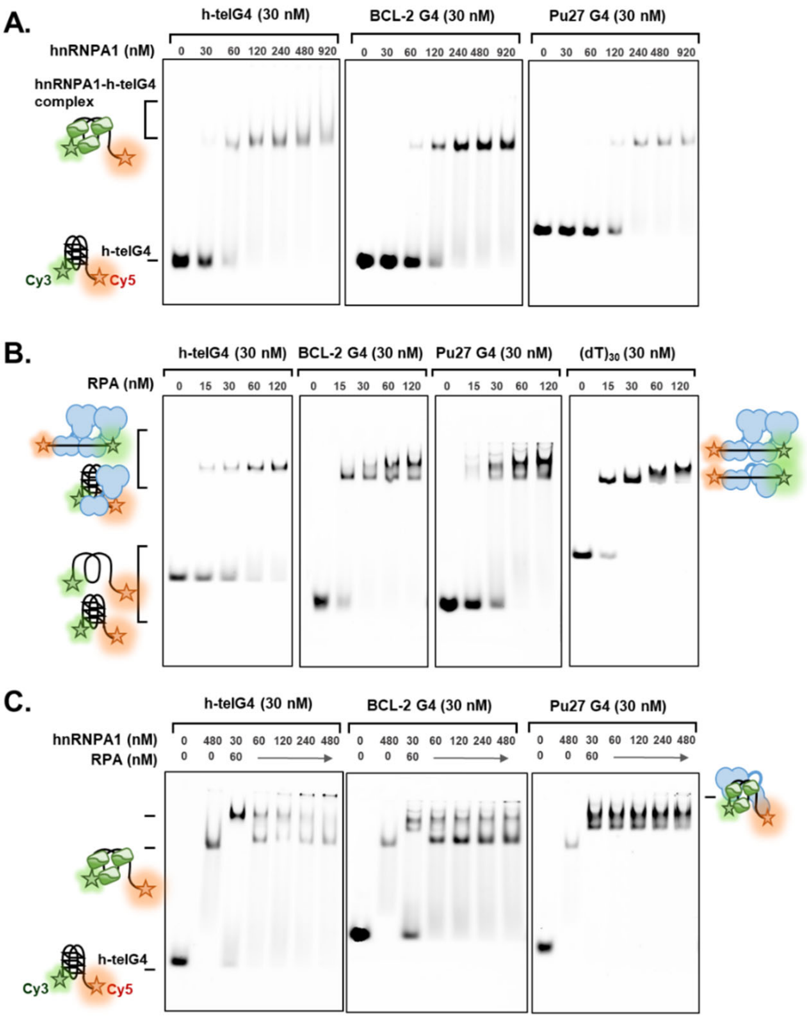


**Supplementary Figure 12. RPA-G4-hnRNPA1 complex is specific to telomeric G-quadruplex. A.** EMSA experiments showing changes in the migration of the fluorescently-labeled DNA (30 nM) in the presence of indicated concentrations of hnRNPA1. Note that hnRNPA1 readily binds two BCL-2 and Pu27 G-quadruplexes, albeit with a lower affinity compared to telomeric G-quadruplex. Note that despite of its ability to bind these G-quadruplexes, no change in the DNA conformation was observed in FRET-based binding experiments (Figure 7A). **B**. EMSA experiments comparing RPA binding to h-telG4, BCL-2G4, Pu27G4 and unstructured ssDNA. In agreement with FRET-based analyses, different bound species and DNA conformations were observed for different DNA substrates. **B**. EMSA experiments showing appearance of the ternary RPA-DNA-hnRNPA1 complex when RPA was bound to telomeric G-quadruplex DNA. Analogous super-shift was observed with other G-quadruplexes only at very high concentrations of hnRNPA1.
